# Substrate recognition and ion coupling mechanism of the human VIAAT

**DOI:** 10.64898/2026.09.12.751155

**Authors:** Yiqing Wei, Kunpeng Ma, Kun Hao, Qinru Bai, Jun Zhao, Xinyu Xie, Renjie Li, Qihao Chen, Yue Li, Hongmei Zhang, Zanxia Cao, Tuo Hu, Yan Zhao

## Abstract

Inhibitory neurotransmission, essential for neural circuit homeostasis and proper neurological function, depends on the efficient sequestration of γ aminobutyric acid (GABA) and glycine into synaptic vesicles. This critical process is mediated exclusively by the vesicular inhibitory amino acid transporter (VIAAT). Despite extensive biochemical and physiological investigation, the molecular mechanism governing VIAAT-mediated transport has remained incompletely understood. Here, we determined structures of human VIAAT in multiple functional states, including apo, GABA-bound, and glycine-bound states, as well as apo state in chloride-free condition. VIAAT adopts a classical LeuT-fold and we elucidated how its large, electronegative binding pocket accommodates both GABA and glycine. Moreover, we resolved two previously unidentified chloride-binding sites. Through integrative molecular dynamics simulations and functional mutagenesis, our results support roles for chloride and the conserved residue E213 in substrate binding and proton-coupled transport. Together, our findings establish a structural framework for VIAAT-mediated inhibitory neurotransmitter transport that illuminates the molecular basis of inhibitory synaptic transmission and provides a foundation for understanding VIAAT dysfunction in epilepsy and related neurodevelopmental disorders.

## Introduction

Inhibitory neurotransmission is essential for counterbalancing excitatory signals in the central nervous system, thereby stabilizing circuit activity and preventing pathological hyperexcitability^1–4^, and also plays a critical role in physiological processes such as motor coordination, sensory processing, and cognitive function^5–7^. γ-Aminobutyric acid (GABA) and glycine are the principal inhibitory neurotransmitters in the mammalian central nervous system, and must be transported into synaptic vesicles via a dedicated transporter to enable normal activity-dependent release^8–11^. Vesicular inhibitory amino acid transporter (VIAAT) is exclusively responsible for loading both GABA and glycine into synaptic vesicles, and its function directly dictates the strength, reliability, and plasticity of inhibitory signaling^12–14^. Disruption of VIAAT function has profound physiological consequences: VIAAT knockout in mice causes embryonic lethality and severe developmental malformations, including omphalocele and cleft palate^15,16^. In humans, missense variants in VIAAT impair GABAergic signaling and underlie a spectrum of genetic epilepsies, ranging from severe developmental and epileptic encephalopathy^17^ to milder genetic epilepsy with febrile seizures plus^18^. These findings underscore the indispensable role of VIAAT in brain development and neural circuit function.

Recent structural studies have advanced our understanding of the transport principles underlying vesicular neurotransmitter transporters, including mechanisms of neurotransmitter recognition and conformational transitions. Examples include detailed investigations of vesicular monoamine transporters (VMATs)^19–23^, vesicular acetylcholine transporter (VAChT)^24,25^, and vesicular glutamate transporters (VGLUTs)^26,27^, all of which belong to the major facilitator superfamily. Despite the central functional role of VIAAT in inhibitory neurotransmission, its architecture has remained experimentally undetermined. Unlike plasma membrane transporters, where GABA and glycine are transported by distinct proteins (GATs for GABA and GlyTs for glycine)^28,29^, VIAAT is responsible for packaging both neurotransmitters into synaptic vesicles^12–14^. The mechanism by which VIAAT recognizes and transports these two different neurotransmitters remains puzzling. Moreover, the driving force underlying VIAAT-mediated transport has also long been controversial. Early studies using isolated synaptic vesicles demonstrated that VIAAT-mediated transport is ATP-dependent and inhibited by the proton gradient-dissipating uncoupler FCCP, suggesting that transport is powered by the electrochemical proton gradient^30,31^. Indeed, VIAAT-mediated transport is associated with proton efflux and vesicle alkalization^32,33^. However, proteoliposome-based experiments reported that valinomycin, which dissipates membrane potential (Δψ), substantially reduced GABA uptake, whereas ammonium sulfate, which selectively dissipates the proton gradient, had only a minor effect. Instead, these studies found that VIAAT-mediated transport strictly requires chloride, with a Hill coefficient of approximately 2.3, leading to the proposal that VIAAT functions as a GABA/2Cl⁻ cotransporter driven by Δψ ^34^. Nevertheless, the precise mechanisms of ion binding and coupling in VIAAT remain unresolved.

To address these mechanistic questions, we determined cryo-electron microscopy structures of human VIAAT in multiple states: apo, GABA-bound, and glycine-bound state in NaCl-containing buffer, as well as apo state in chloride-free phosphate buffer. By integrating molecular dynamics simulations and neuron-based electrophysiological studies, our study provides mechanistic insights and a structural framework for understanding the molecular mechanisms underlying substrate recognition, ion coupling, and conformational transitions during the transport cycle. Collectively, these findings establish a long-sought structural framework for understanding VIAAT function.

## Results

### Structure determination and architecture of the human VIAAT

To elucidate the molecular mechanisms of VIAAT, we generated a wild-type VIAAT construct harboring a C-terminal Twin-Strep tag for purification, along with an IRES-mCherry cassette to monitor expression. To validate the activity of this construct, we established a functional assay using whole-cell patch-clamp recordings in cultured hippocampal neurons (Fig. 1a). Knockdown of endogenous VIAAT with short hairpin RNA (shRNA) targeting the endogenous mouse *VIAAT* transcript (shRNA^VIAAT^) abolished inhibitory postsynaptic currents (IPSCs), whereas exogenous expression of the Twin-Strep–tagged VIAAT, with optimized codons rendering it resistant to shRNA^VIAAT^, efficiently restored synaptic transmission (Fig. 1b), confirming that the tagged protein retains its physiological function. This construct was subsequently used for protein expression and cryo-EM sample preparation. Purification using affinity and size-exclusion chromatography yielded a symmetric peak (Extended Data Fig. 1a), indicating high monodispersity and suitability for structural studies. Coomassie-stained SDS–PAGE of the peak fraction revealed multiple bands between 42 and 72 kDa (Extended Data Fig. 1b), a phenomenon often observed for membrane proteins, which may arise from heterogeneity such as post translational modifications. Using single-particle cryo-EM, we determined the structures of VIAAT in the apo state and in the presence of the substrate GABA or glycine, at resolutions of 3.3 Å, 2.7 Å and 3.1 Å, respectively (Fig. 1c and Extended Data Fig. 2-4). We also determined the apo structure of VIAAT in chloride-free phosphate buffer, achieving a resolution of 3.0 Å (Extended Data Fig. 5). All maps exhibit well-defined features, including clear side-chain densities, allowing us to unambiguously build the atomic models for each structure (Extended Data Fig. 2-5 and Extended Data Table 1).

**Fig. 1.**
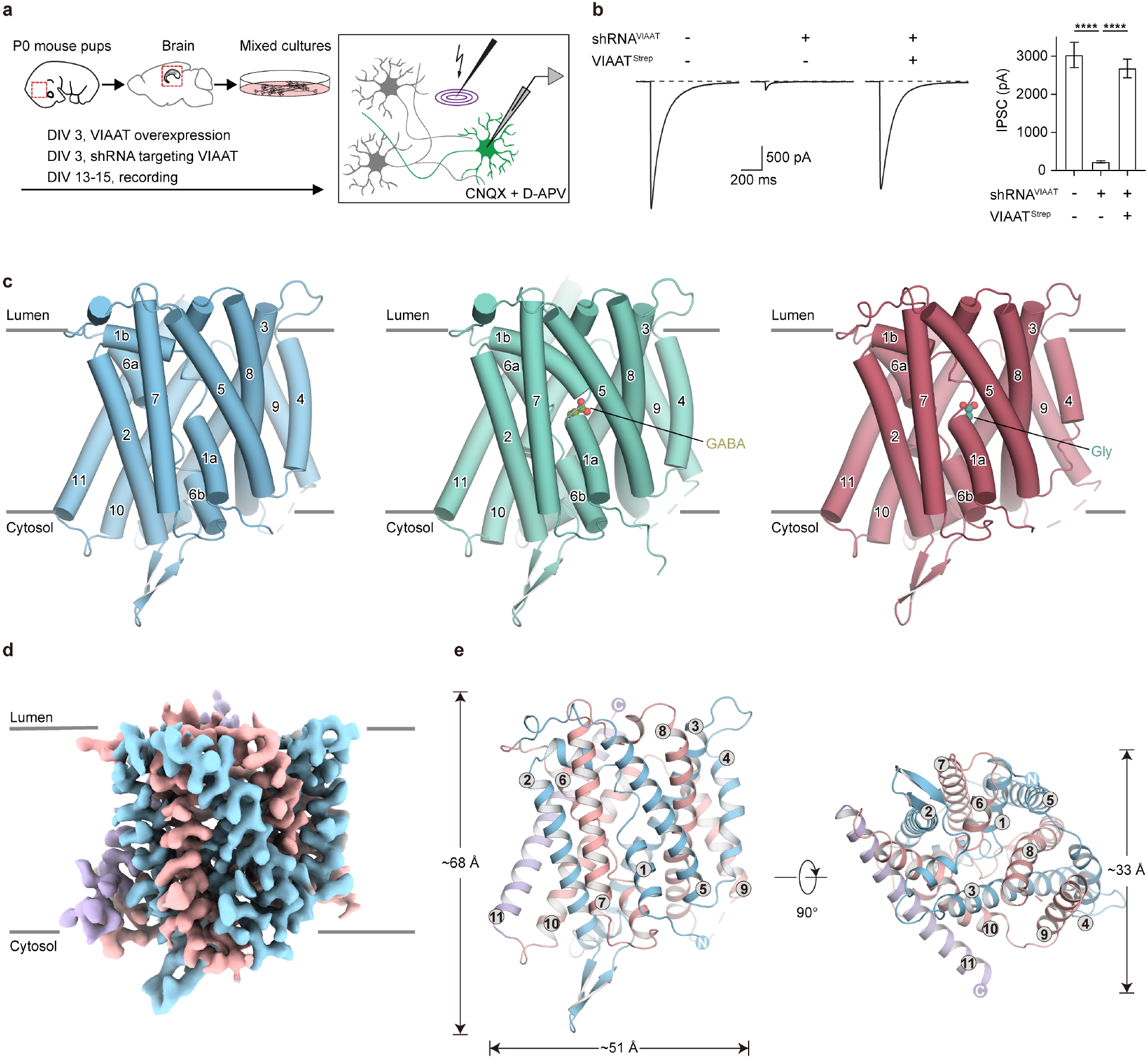
Functional characterization and architecture of human VIAAT. **a**, Schematic of the electrophysiological recording paradigm. Evoked inhibitory postsynaptic currents (IPSCs) were recorded from cultured hippocampal neurons. Primary neurons were transduced with shRNA or rescue lentiviruses at DIV3 and recordings were performed at DIV13–15 (see Methods). **b**, Left, representative traces of evoked IPSCs recorded from control (shRNA) and VIAAT-rescue neurons. Right, quantification of IPSC amplitudes under the indicated conditions. Data are shown as mean ± s.e.m.; statistical significance was assessed using unpaired t-test; ****P < 0.0001. **c**, Cartoon representations of the VIAAT structures in the apo, GABA-bound, and glycine-bound states. GABA and glycine are shown as olive and teal spheres, respectively. Nitrogen and oxygen atoms are coloured blue and red, respectively. Transmembrane helices (TM1–TM11) are labelled, and the lumenal and cytosolic sides are indicated. **d**, Cryo-EM density map of VIAAT reconstituted in nanodiscs and captured in a lumen-facing open conformation. Densities corresponding to TM1–5, TM6–10, TM11, and lipid molecules are coloured light blue, light pink, lavender, and yellow, respectively. The lumenal and cytosolic sides are indicated. **e**, Cartoon representations of the VIAAT apo model shown in side and top-down views. TM1–5, TM6–10, and TM11 are coloured as in D. Overall dimensions as well as transmembrane helices and loops are indicated.

Human VIAAT consists of 525 amino acids, of which residues 113-521 were resolved in our structures. The structure reveals an 11-transmembrane (TM) architecture, measuring 68 Å in height, 51 Å in width (parallel to the membrane plane), and 33 Å in depth (perpendicular to the membrane plane) (Fig. 1d, e). VIAAT adopts a compact, pseudo-symmetric fold composed of two inverted repeats (TMs 1–5 and TMs 6–10), flanked by an additional peripheral helix (TM11). Both TM1 and TM6 contain an unwound segment, dividing each into two short helices. This organization represents the classical LeuT-fold architecture (Fig. 1e and Extended Data Fig. 6a). The apo structure of VIAAT (VIAAT^Apo^) reveals a large cavity open toward the luminal side, while the cytosolic gate remains sealed, representing a lumen-facing open conformation (Extended Data Fig. 6b).

Structural superimposition of VIAAT^Apo^ onto outward-facing open GlyT1, using the scaffold domain (TMs 3, 4, 8, and 9) as a reference, shows that VIAAT adopts the canonical SLC6 fold but also displays several unique features (Extended Data Fig. 6c-e). Most transmembrane helices exhibit moderate displacements relative to GlyT1 (Extended Data Fig. 6c); however, pronounced differences arise in EL4, TM6a, and TM10. In VIAAT, EL4 is markedly shorter than in GlyT1 (Extended Data Fig. 6d). Whereas the EL4 in GlyT1 forms an extracellular lid over the central cavity, the shorter EL4 in VIAAT leaves the luminal vestibule more exposed and generates additional space above the substrate-binding pocket. Moreover, VIAAT contains a pronounced π-helical bulge in the central portion of TM10, causing the helix to bend and shift toward TM6 relative to its position in GlyT1 (Extended Data Fig. 6e). This displacement reduces the lateral space available on the side of TM6a. Consistent with this altered geometry, TM6a in VIAAT adopts a more closed conformation, tilting by roughly 30° toward the central axis compared with the more open TM6a conformation in GlyT1 (Extended Data Fig. 6e). This rearrangement likely accommodates the shorter EL4 and the displacement of TM10 in VIAAT. On the cytosolic side, VIAAT features a prominent loop between TM2 and TM3 that folds into a short antiparallel β-sheet (Extended Data Fig. 6f). This loop is substantially larger than the corresponding region in GlyT1 and is enriched in charged residues—including aspartate, glutamate, and arginine—representing a unique structural element of VIAAT (Extended Data Fig. 6f).

An increasing number of missense variants in VIAAT have been implicated in genetic epilepsy, including developmental and epileptic encephalopathy and genetic epilepsy with febrile seizures plus^17,18^. Our structural data provide a precise structural framework for interpreting the functional consequences of these pathogenic mutations (Extended Data Fig. 6g and Extended Data Table 2). For instance, F322^TM^^6^ is located within the substrate-binding pocket, with its side chain positioned approximately 4.8 Å from the bound GABA in VIAAT^GABA^. The F322C mutation is likely to interfere with substrate coordination (Extended Data Fig. 6h). P395^TM8^ and L468^TM10^ are situated within the central regions of their respective transmembrane helices (Extended Data Fig. 6i). Pathogenic mutations such as P395L or L468P are expected to substantially alter helix geometry, potentially impairing transport function. Furthermore, mutations G461D, T464R, and G465S are located at TM10, which has been implicated in regulating extracellular gate dynamics in LeuT-fold transporters^35^. These mutations introduce bulkier side chains, which may destabilize extracellular gate closure and disrupt conformational transition (Extended Data Fig. 6j). Consistent with our structural analysis, functional analysis of the disease-associated variants F322C, P395L, L468P, G461D, T464R, and G465S showed that all six mutants displayed markedly reduced transport activity (Extended Data Fig. 6k).

### Substrate recognition of VIAAT

VIAAT transports the inhibitory neurotransmitters glycine and GABA into synaptic vesicles, with a reported *K*_m_ of 5 mM for GABA. Glycine, which has a lower affinity for VIAAT, competes with GABA for transport and exhibits an IC50 of 27.5 mM^13^. To uncover the molecular mechanism underpinning substrate transport, we determined the structure of VIAAT in the presence of high concentration of GABA (VIAAT^GABA^) and glycine (VIAAT^Gly^), respectively. In both complexes, we observed a well-defined rod-shaped density within the central cavity, which is absent from the equivalent position in the structure of VIAAT^Apo^ and thus may represent a bound substrate (Fig. 2a). In the structure of VIAAT^GABA^, the overall structure is well aligned with VIAAT^Apo^ (Extended Data Fig. 7a), indicating it adopts a lumen-facing conformation, yet the substrate-binding pocket is fully shielded from solvent on both luminal and cytosolic sides, implying that the structure is captured in a lumen-facing occluded state (Fig. 2b). The binding pocket forms a strongly electronegative environment (Fig. 2b). The carboxyl group of GABA is oriented toward the lumen, while its amino group points toward the cytosol (Fig. 2b, c), interacting with surrounding residues and water molecules (Fig. 2d). Specifically, the carboxyl group forms hydrogen bonds with the hydroxyl group of Y221^TM3^ and the backbone amides of M132^TM1^ and F133^TM1^, as well as a water-mediated interactions with the backbone amide of G131^TM1^ and the side chains of S316^TM6^ and K351^TM7^. The amino group hydrogen-bonds with the backbone carbonyls of A128^TM1^ and T318^TM6^ and the side-chain hydroxyl of T318^TM6^. Additional stabilization is also provided by water-mediated contacts involving N127^TM1^, E213^TM3^ and T217^TM3^. The carbon skeleton of GABA is accommodated by hydrophobic residues, including F133^TM1^ and F315^TM6^ (Fig. 2c, d). In the VIAAT^Gly^ structure, the carboxyl group of glycine forms hydrogen bonds with the backbone amide of G131^TM1^ and the phenolic hydroxyl of Y221^TM3^, while the amino group interacts with the backbone carbonyl of A128^TM1^ and the side chain of T318^TM6^. Several ordered water molecules surround the amino group of glycine, forming an extended hydrogen-bonding network that indirectly engages N127^TM1^, E213^TM3^, and T217^TM3^, further stabilizing the glycine binding (Fig. 2e).

**Fig. 2.**
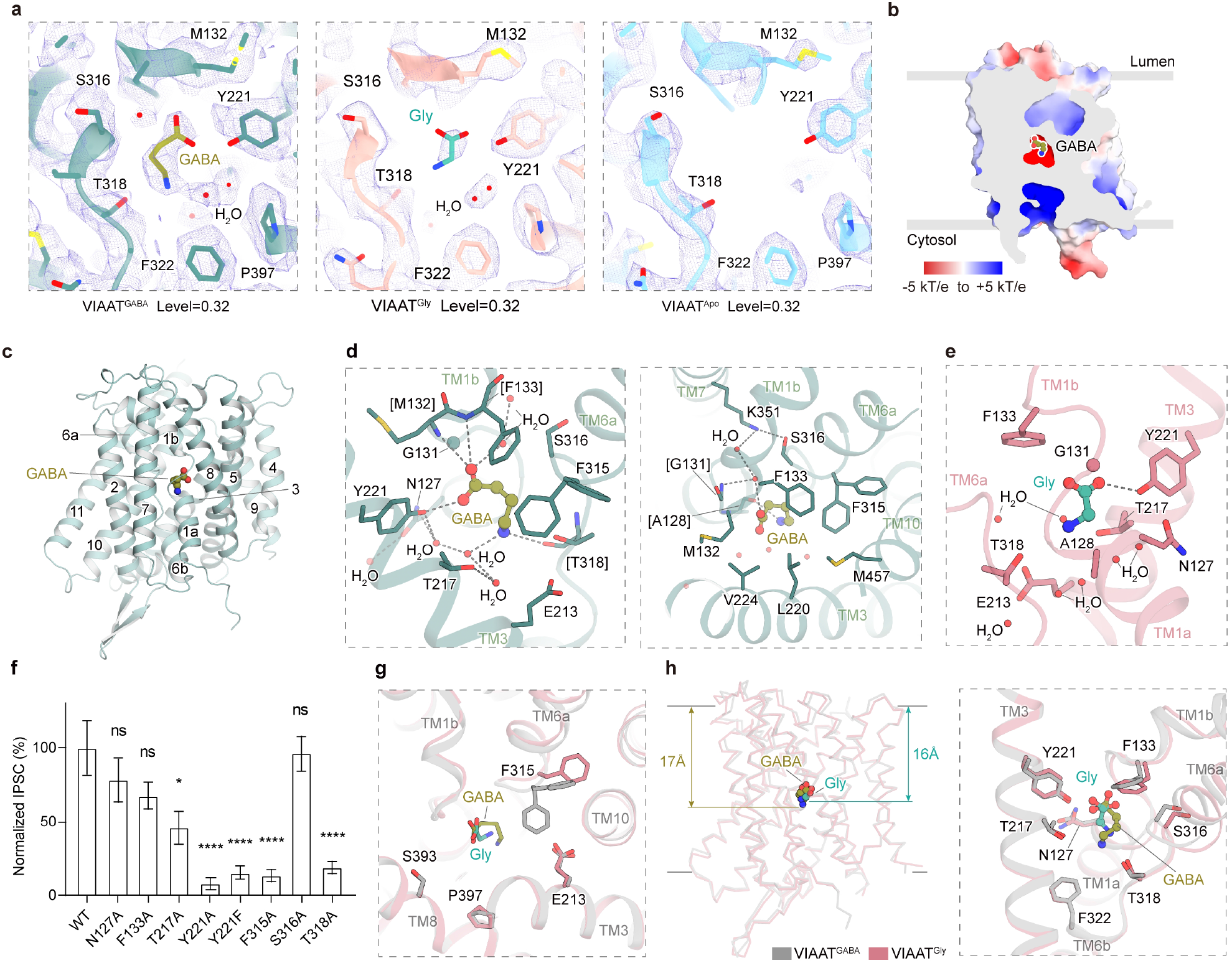
GABA and glycine recognition by VIAAT. **a**, Comparison of cryo-EM densities within the substrate-binding pocket of VIAAT in the GABA-bound (left), glycine-bound (middle), and apo (right) states. Densities contoured at 0.32 are shown as violet mesh. Residues lining the substrate-binding pocket are depicted as sticks. GABA and glycine are shown as olive and teal sticks, respectively. Nitrogen, oxygen, and sulfur atoms are coloured blue, red, and yellow, respectively. **b**, Electrostatic surface potential of VIAAT^GABA^, shown as a slice through the substrate-binding pocket. GABA is displayed as spheres. Positively and negatively charged surfaces are coloured blue and red, respectively. **c**, Overall structure of VIAAT in the GABA-bound state shown in cartoon representation. GABA is rendered as spheres and labelled. Transmembrane helices are labelled. **d,e,** Close-up views of the substrate-binding pocket showing interactions with GABA (d) and glycine (e), respectively. Substrates are depicted in olive and cyan ball-and-stick representations. Key interacting residues are shown as sticks and labelled. Water molecules are shown as red spheres. Dotted lines indicate potential hydrogen-bond interactions. **f**, Normalized IPSC amplitudes for wild-type (WT) VIAAT and substrate-binding mutants. IPSC amplitudes of mutants were normalized to those of WT. Data are presented as mean ± s.e.m. from biologically independent experiments (n = 5–18). Statistical significance was assessed using an unpaired t-test with α = 0.05. *P < 0.05; **P < 0.01; ***P < 0.001; ****P < 0.0001. **g**, Comparison of the conformations of F315 in VIAAT^GABA^ (grey) and VIAAT^Gly^ (light pink). F315 adopts two different conformations in the VIAAT^GABA^ structure. **h**, Overview (left) and close-up view (right) of the superposition of the substrate-binding pockets of VIAAT in the VIAAT^GABA^ and VIAAT^Gly^. Key residues are shown as sticks.

To assess the functional relevance of these interactions, we performed mutagenesis on key binding residues, including residues N127^TM1^, F133^TM1^, T217^TM3^, Y221^TM3^, and T318^TM6^ (Fig. 2f). Mutations Y221F/A and T318A nearly abolished transport activity, while N127A, F133A and T217A resulted in moderate reductions (Fig. 2f). After train stimulation-induced depletion of inhibitory synaptic vesicles, the N127A and F133A mutants exhibited a significantly slower rate of IPSC recovery compared to WT (Extended Data Fig. 8a). All mutants analyzed in this study showed robust expression and synaptic localization comparable to WT (Extended Data Fig. 8b, c), and the functional data were normalized to expression levels. Additionally, in the substrate-bound structures, we observed that residue F315^TM6^ adopts distinct conformations. In the VIAAT^GABA^ structure, F315^TM6^ adopts two alternative rotamers: in rotamer 1 (rot1), the side chain swings away to the lumen side and toward TM10, whereas in rotamer 2 (rot2), it points toward TM3 and contribute to GABA binding (Fig. 2g). By contrast, in the VIAAT^Gly^ structure, F315^TM6^ predominantly adopts rot1 (Fig. 2g). The rot1 allows forming a continuous path to the lumen side, representing a lumen-facing open state (Extended Data Fig. 7b), while rot2 seals the cavity, leading to an occluded state (Fig. 2b and Extended Data Fig. 7c). Substitution of F315^TM6^ with alanine almost eliminated transport activity (Fig. 2f), highlighting the critical role of this hydrophobic residue in mediating conformational transitions during substrate transport.

Structural comparison of VIAAT^GABA^ and VIAAT^Gly^ reveals that both substrates engage the binding site in nearly identical orientations and most of the residues involved in substrate recognition adopt similar conformations (Fig. 2h). However, due to its longer carbon backbone, GABA extends deeper into the binding cavity than glycine (Fig. 2h). The substrate is in close proximity to the unwound regions of TM1 and TM6 in VIAAT, consistent with previous studies suggesting that these regions in the LeuT family are important for substrate recognition^36–39^. Although VIAAT, GlyT1, GlyT2, and GAT1 all bind substrate within this conserved region (Extended Data Fig. 7d), their binding affinities differ markedly. The substrate-binding cavity in VIAAT is approximately 323 Å^3^—substantially larger than either GABA or glycine—whereas GlyT1, GlyT2, and GAT1 possess cavities of approximately 122, 89, and 178 Å^3^, respectively, volumes that more tightly match their substrates. The larger substrate-binding pocket likely contributes to the lower affinity of VIAAT for both neurotransmitters. In addition to its canonical substrates GABA and glycine, previous studies have reported that β-alanine and γ-vinyl-GABA were competitive inhibitors of GABA transport through VIAAT ^40,41^. These observations demonstrate that the large substrate-binding cavity of VIAAT possesses high ligand-recognition plasticity and is capable of accommodating structurally related substrate analogs.

### Ion coupling during the substrate translocation of VIAAT

The ions coupled to transport by vesicular neurotransmitter transporters have been extensively investigated. VMATs utilize the vesicular proton gradient, consuming two protons per transport cycle^42,43^. The VGLUTs depend on both the membrane potential and the vesicular proton electrochemical gradient to drive glutamate uptake^44–47^. However, the ion-coupling mechanism underlying VIAAT function remains incompletely understood. Previous studies have demonstrated that both protons and chloride ions contribute to VIAAT activity^30–34^. The samples of VIAAT^Apo^, VIAAT^GABA^, and VIAAT^Gly^ were prepared in NaCl buffer. To investigate whether chloride may interact with VIAAT, we prepared VIAAT samples in a chloride-free phosphate buffer (VIAAT^PO4^) and determined its structure at 3.0 Å resolution (Extended Data Fig. 5 and Extended Data Table 1). After careful inspection of the cryo-EM maps of VIAAT^GABA^ and VIAAT^PO4^, we identified two putative chloride-binding sites based on characteristic coordination geometry and local density features (Fig. 3a). The putative Cl1 site is located near N127^TM1^, Q130^TM1^, T271^TM5^, H274^TM5^, and K389^TM8^ (Fig. 3b), whereas the putative Cl2 site is defined by S319^TM6^, Q320^TM6^, and H344^TM7^ (Fig. 3c). At these two positions, we observed distinct sphere-shaped densities in the VIAAT^GABA^ map, but these were absent in VIAAT^PO4^ (Fig. 3d, e). Molecular dynamics simulations showed persistent chloride occupancy at these sites over 400 ns, providing additional support for their assignment (Extended Data Fig. 9a-c).

**Fig. 3.**
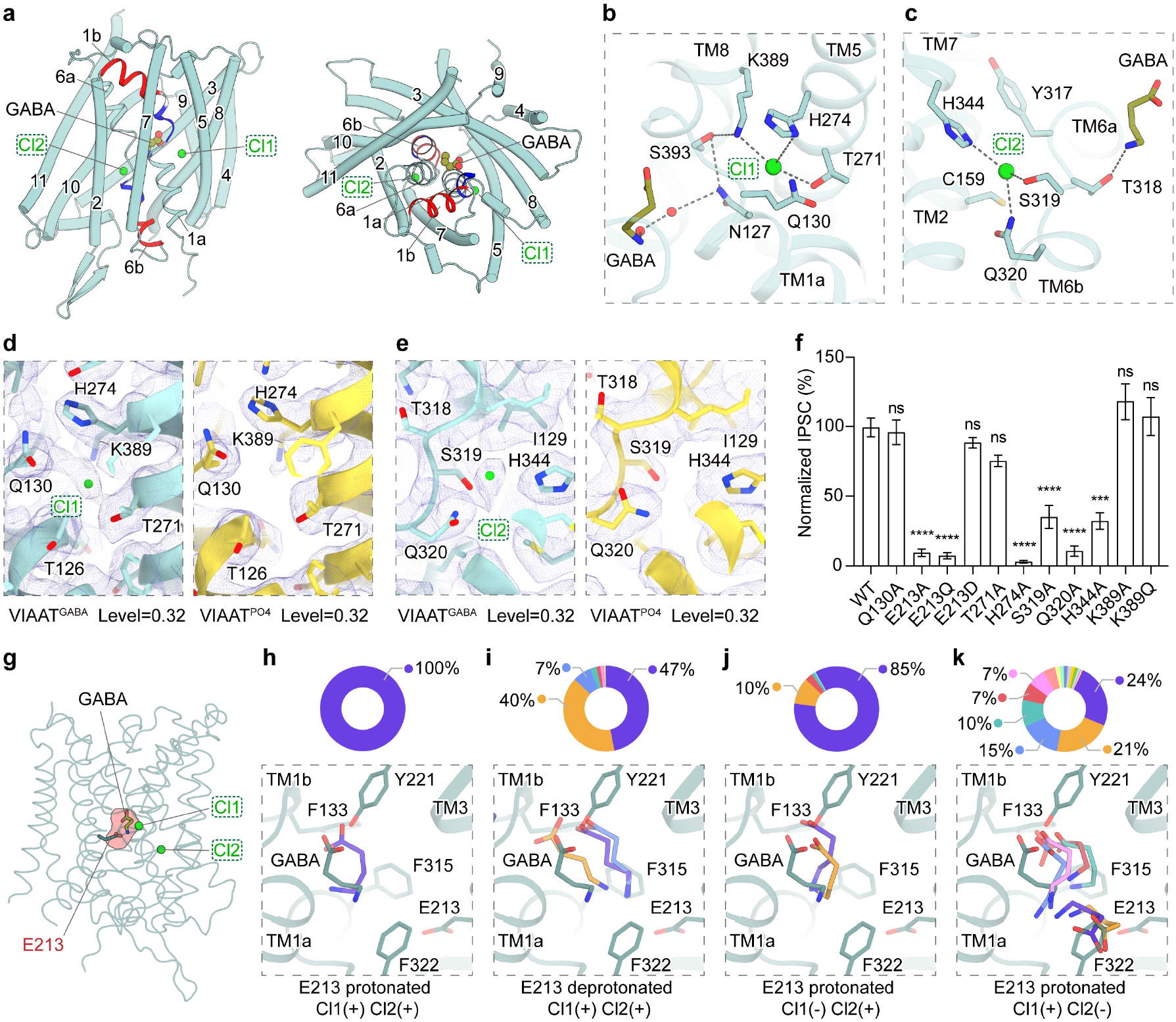
Chloride ions and proton coupling in VIAAT. **a**, Side view (left) and top-down view (right) of the overall structure of VIAAT in the GABA-bound state, with putative chloride ions highlighted. Chloride ions and GABA are shown as green and olive spheres, respectively. TM1 and TM6 are shown as cartoons, whereas the remaining transmembrane helices are shown as cylinders. TM1b and TM6a are coloured by dipole potential, with blue and red indicating positive and negative potentials, respectively. **b,c,** Close-up views of the Cl1 (b) and Cl2 (c) binding pockets. Key residues involved in ion coordination are shown as sticks. Potential hydrogen-bond interactions are indicated by black dotted lines. **d,e,** Comparison of cryo-EM densities at the Cl1 (d) and Cl2 (e) sites in the GABA-bound (left) and chloride-free apo (right) states of VIAAT. Densities contoured at 0.32 are shown as violet mesh, with the corresponding models displayed as cartoons and sticks. Chloride ions are depicted as green spheres. **f**, Normalized IPSC amplitudes for wild-type (WT) VIAAT and ion-binding mutants. IPSC amplitudes of mutants were normalized to those of WT. Data are presented as mean ± s.e.m. from biologically independent experiments (n = 5–18). Statistical significance was assessed using an unpaired t-test with α = 0.05. *P < 0.05; **P < 0.01; ***P < 0.001; ****P < 0.0001. **g,** Overview of VIAAT in the GABA-bound state highlighting the titratable residue E213 located in the central pocket. The chloride ions are shown as green spheres. The GABA-binding pocket is outlined in red, and GABA is rendered as olive sticks. **h-k,** Frame clustering of molecular dynamics simulations starting from the cryo-EM VIAAT^GABA^ structure under different conditions. Clusters were obtained from three independent 400 ns trajectories using a 1.5 Å RMSD cut-off calculated for GABA and surrounding residues within 5 Å for each condition. Top: Each cluster is shown in a distinct color on the ring charts. Dominant clusters exceeding a 5% population threshold are labeled. Bottom: Superposition of VIAAT^GABA^ (deep green) and dominant clustered structures (colored as in the ring charts).

These putative chloride-binding sites were further compared with the sodium-binding sites of LeuT-fold transporters, using GlyT1 as a representative example (Extended Data Fig. 10a). We found that the residues coordinating sodium and chloride in GlyT1 are not conserved in VIAAT, and conversely the chloride coordinating residues in VIAAT are not conserved in GlyT1 (Extended Data Fig. 10b, c), highlighting the distinct structural basis underlying their specific ion selectivity. Moreover, we observed that the putative Cl1 site in VIAAT roughly overlaps with the Na2 site in GlyT1 (Extended Data Fig. 10b). The Na2 site has been shown to play a pivotal functional role in LeuT-fold transporters. Previous findings revealed that sodium binding at this position stabilizes the outward-facing conformation^48–50^. We therefore propose that the chloride ion coordinated at the Cl1 site in VIAAT may fulfill a similar structural function. We also introduced mutations at these positions to further assess the functional importance of the chloride-coordinating residues and found that H274A, S319A, Q320A, and H344A did not significantly affect protein expression, but markedly impaired transport activity (Fig. 3f), highlighting the essential role of these residues in transporter function. Although our structural, mutant functional assays and MD analyses support the identification of these putative chloride-binding sites in VIAAT and suggest a possible role for chloride, the physiological relevance of these putative chloride-binding sites requires further validation.

To identify residues potentially involved in proton-coupled transport, we focused on titratable residues within the highly electronegative substrate-binding pocket. Among residues in this pocket, only E213^TM3^ is titratable and is highly conserved across VIAAT orthologs from different species (Fig. 3g, Extended Data Fig. 11). The predicted pKa in the lumen-facing open VIAAT^Apo^ structure of E213^TM3^ is 6.59, calculated using PropKa program^51^. Considering that the physiological pH in the vesicular lumen is approximately 6.2-6.4^33^, we propose that E213^TM3^ is likely to be protonated on the luminal side. To further evaluate the functional role of E213^TM3^, we generated a cytosol-facing homology model of VIAAT (VIAAT^Cyto^) using the inward-facing structure of SLC38A9 as a template^52^, given the close homology between VIAAT and SLC38A9. In this model, E213^TM3^ is accessible from the cytoplasmic side (Extended Data Fig. 10d), and its predicted pKa is 6.67, indicating it would tend to be deprotonated at the cytosolic pH of ∼7.4. These observations suggest that E213 can be alternately exposed to the luminal or cytoplasmic side during the transport cycle and may undergo protonation and deprotonation, respectively. Both E213A and E213Q abolished transport activity, whereas E213D, which retains a titratable carboxyl group, exhibited transport activity comparable to that of the wild type (Fig. 3f), supporting potential role of E213 in proton coupling.

To further investigate the ion-coupling mechanism of VIAAT, we performed all-atom molecular dynamics simulations under various ionic conditions. For each condition, three independent trajectories of 400 ns were conducted. Throughout all simulations, the structure of VIAAT is very stable with low RMSD value of backbone (Extended Data Fig. 9a). To validate the effects of different conditions on GABA and ion binding, we first evaluated the impact of E213 protonation on substrate binding by performing clustering analysis on GABA and residues within 5 Å of the ligand, using an RMSD cutoff of 1.5 Å. When E213 was protonated, only a single dominant conformation of substrate GABA was identified, closely resembling the GABA pose observed in the VIAAT^GABA^ structure (Fig. 3h). In contrast, deprotonation of E213 resulted in multiple distinct clusters with considerable conformational variation from the binding pose determined in the VIAAT^GABA^ structure (Fig. 3i). Importantly, we found that the amino group of GABA was displaced toward the carboxyl group of E213 (Fig. 3i), potentially establishing a stronger electrostatic interaction between these two groups compared to the protonated state of E213. This observation is consistent with findings in other proton-dependent antiporters, where protonation and deprotonation of key acidic residues within the substrate-binding pocket may modulate the substrate release process^19,53^.

Moreover, we systematically compared the effects of the presence or absence of either or both chloride ions on substrate binding, to evaluate the potential functional role of each chloride ion. Both Cl1 and Cl2 ions remained highly stable throughout each trajectory, regardless of the protonation state of E213 (Extended Data Fig. 9b, c). Under protonated E213 conditions, removal of Cl1 from the starting model resulted in Cl2 remaining stably coordinated (Extended Data Fig. 9c), while GABA adopted one predominant pose similar to that in the starting model (Fig. 3j). In contrast, simulations initiated from a Cl2-depleted state revealed pronounced instability in GABA binding (Extended Data Fig. 9d); under these conditions, GABA sampled many distinct poses, indicating substantial conformational variability (Fig. 3k). Cl1 also deviated from its initial binding site (Extended Data Fig. 9b). These simulations suggest that Cl1 may not be essential for substrate association. Instead, it occupies a position similar to Na2 in other LeuT-fold transporters, likely playing a comparable role in facilitating conformational transitions^48–50,54^. Conversely, Cl2 appears to be critical for substrate binding, as its depletion significantly destabilizes GABA binding.

### Conformational transition mechanism of VIAAT

Secondary active transporters operate through conformational transitions among outward-facing, occluded, and inward-facing states^55^. In the LeuT-fold transporters, specific intramolecular interactions stabilize distinct conformations along the transport cycle. In GlyT1, for example, the R71–D474 interaction is critical for maintaining the inward-facing conformation, whereas on the cytoplasmic side, a RxxW motif in the N-terminal tail engages F55, Y331, and D432 to stabilize the outward-facing state^37^. These interactions are highly conserved across the SLC6 family^37,56^. Although VIAAT also adopts a LeuT fold, most of these hallmark interactions are not preserved, suggesting that VIAAT employ distinct structural determinants to regulate its conformational equilibrium.

In the lumen-facing VIAAT^Apo^ structure, TM1a is locked through interactions with surrounding structural elements. For instance, K113^IL1^ on the N-terminal loop forms a salt bridge with E403^TM8^, and E120^TM1^ engages in electrostatic interactions with K261^TM5^ and K265^TM5^ (Fig. 4a-c). These contacts hold TM1 close to the scaffold domain and prevent its dilation, a movement necessary for transition to the cytosol-facing conformation^37^. Structural comparison between VIAAT^Apo^ and VIAAT^Cyto^ revealed that the scaffold domain (TM3, TM4, TM8, and TM9) remains largely rigid during the conformational transition. In contrast, TM1b and TM6a rotate by approximately 32° and 19°, respectively, whereas TM1a and TM6b swing outward by approximately 34° and 45°, respectively, converting the central cavity from a lumen-facing to a cytosol-facing configuration (Extended Data Fig. 12a). Accompanying these movement, TM2 and TM7 tilt toward the scaffold domain by 18° and 12°, while extracellular loops EL3 and EL4 shift by approximately 7.5 Å and 7.3 Å towards membrane plane, respectively (Extended Data Fig. 12a, b). In the VIAAT^Cyto^ model, residues including Y139^TM1b^, H143^TM1b^, K305^TM6^, E368^EL4^ and H452^EL5^ converge to form a compact electrostatic and hydrogen-bonding network that is absent in lumen-facing VIAAT^Apo^ (Fig. 4d, e), potentially stabilizing the cytosol-facing conformation of VIAAT. Structural comparison and sequence alignment demonstrated that these residues are not conserved in the SLC6 family but are highly conserved among VIAAT homologs from different species (Extended Data Figs. 11 and 12e, f).

**Fig. 4.**
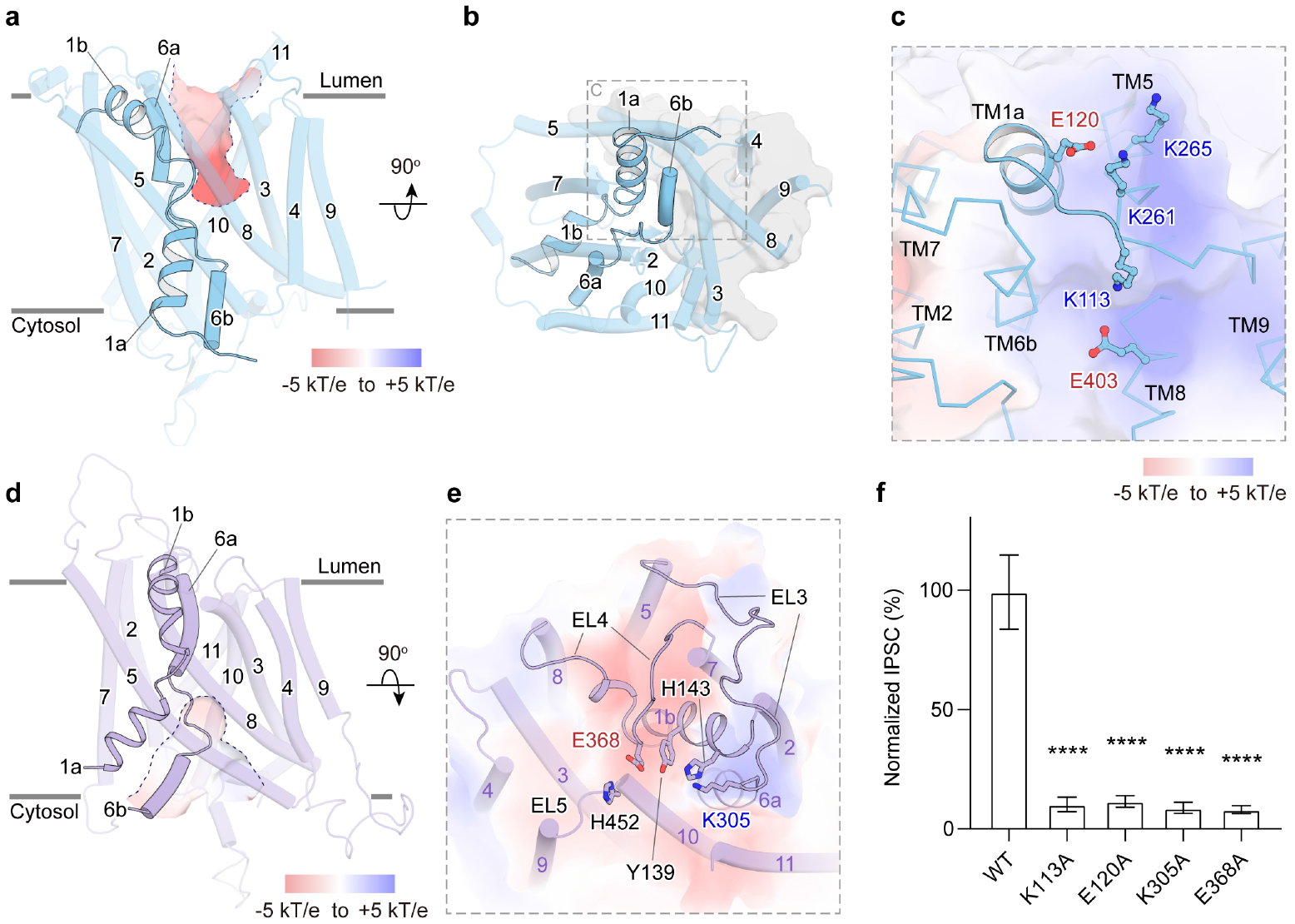
Interactions governing conformational transition of VIAAT. **a,b,** Side view (a) and top-down view (b) of the overall structure of VIAAT in the apo state captured in a lumen-facing conformation. TM1a is shown as a cartoon, whereas the remaining transmembrane helices are shown as cylinders. The electrostatic surface potential along the pathway from the substrate-binding pocket to the lumen is displayed and outlined by a dotted line. **c,** Close-up view of key residues stabilizing the lumen-facing state of VIAAT. Critical residues are depicted as ball-and-stick models, and the electrostatic surface potential is shown. **d**, Overall structure of a homology model of VIAAT in the cytosol-facing state, with TM1a shown as a ribbon. The electrostatic surface potential along the pathway from the substrate-binding pocket to the cytosol is shown and outlined by a dotted line. **e**, Top-down view of homology model of VIAAT in the cytosol-facing state. Potential key residues stabilizing cytosol-facing state are shown as sticks. The electrostatic surface potential is shown. **f,** Normalized IPSC amplitudes for wild-type (WT) VIAAT and gating-related mutants. IPSC amplitudes of mutants were normalized to those of WT. Data are presented as mean ± s.e.m. from biologically independent experiments (n = 6–7). Statistical significance was assessed using an unpaired t-test with α = 0.05. *P < 0.05; **P < 0.01; ***P < 0.001; ****P < 0.0001.

To assess the functional relevance of these interactions, we performed mutagenesis studies. Substitution of K113^IL1^, E120^TM1^, K305^TM6^, or E368^EL4^ with alanine severely impaired or abolished transport activity (Fig. 4f). These findings support their important role for VIAAT function. We propose that disrupting these interactions perturbs the conformational balance of VIAAT, leading to over-stabilization of particular states and impaired transport activity.

## Discussion

In this study, we developed a robust functional assay using cultured neurons to record inhibitory postsynaptic currents, enabling us to directly assess VIAAT activity in a physiologically relevant context. Using single particle cryo-EM, we determined the structures of human VIAAT in its apo state and in complexes with either GABA or glycine. Structural analysis revealed that both the apo and glycine-bound forms adopt a lumen-facing open conformation, whereas the GABA-bound structure is captured in a lumen-facing occluded state. Notably, the side chain of F315 adopts distinct rotamers in these states. In the occluded conformation, F315 rotates to seal the extracellular vestibule, while in the open conformation it points toward TM10 allowing access to the binding pocket. Functional assays demonstrated that F315 is essential for transport activity, likely due to its pivotal role in substrate binding and conformational transitions. Both GABA and glycine are accommodated within a highly electronegative binding pocket and occupy similar positions, highlighting a shared substrate recognition mechanism despite their chemical differences. Our structural analyses further reveal that, although the overall pattern of conformational transitions in VIAAT is highly conserved among LeuT-fold transporters, the unique structural motifs and interaction networks that stabilize specific conformational states are conserved within VIAAT homologs but not in the SLC6 family.

The ion coupling mechanism of VIAAT has long been debated, with both proton and chloride having been proposed to be important for transport activity^30–32^. However, the molecular basis underlying this process remains incompletely understood. In this study, we identified two putative chloride-binding sites in VIAAT, with the putative Cl1 site located near N127^TM1^, Q130^TM1^, T271^TM5^, H274^TM5^, and K389^TM8^ and the putative Cl2 site defined by S319^TM6^, Q320^TM6^, and H344^TM7^. Structural analysis revealed that these chloride sites are distinct from the sodium-binding sites found in SLC6 family transporters. Furthermore, mutagenesis of key coordinating residues supported their functional importance. Notably, the putative Cl1 site is positioned similarly to the Na2 site in SLC6 transporters, underscoring both evolutionary divergence and conservation among LeuT fold transporters in their selection of coupling ions. In addition, our integrative approach combining molecular dynamics simulations with functional assays supports the conserved E213 residue as a critical determinant of proton-coupled transport. The protonation state of E213 may modulate the energy barrier for substrate dissociation, providing a mechanistic explanation for the observed coupling of proton flux to inhibitory neurotransmitter transport. While our structural analyses, mutant functional assays, and MD simulations support the identification of the putative binding sites for chloride and proton ions, additional experimental data, such as direct measurements of substrate affinity and ion gradient-controlled reconstitution experiments, would provide valuable complementary insights into the specific roles of these ions in transporter activity. Collectively, our findings establish a structural and mechanistic blueprint for VIAAT-mediated inhibitory neurotransmitter transport. The elucidation of substrate recognition, ion coupling, and conformational dynamics advances our fundamental understanding of inhibitory synaptic transmission, and provides a foundation for understanding VIAAT dysfunction in epilepsy and related neurodevelopmental disorders.

## Methods

### Clone, Expression and Purification of human VIAAT

The full-length wild-type human vesicular GABA transporter (VIAAT, SLC32A1; UniProt Q9H598) fused to a C-terminal twin-strep tag and an IRES–mCherry cassette was cloned into the pEG BacMam vector. The IRES sequence enabled independent translation of VIAAT and the fluorescent reporter. The verified construct was transformed into DH10Bac *E. coli* for bacmid generation. Positive white colonies were selected for bacmid extraction, which was then used to transfect Sf9 cells for production of recombinant baculovirus. For protein expression, HEK293F cells at a density of approximately 2×10^6^ cells per mL were infected with 1% v/v P2 baculovirus and cultured at 37 °C under 5% CO_2_. To enhance protein expression, sodium butyrate was added 24 h after transduction to a final concentration of 10 mM, after which the temperature was reduced to 30 °C while maintaining 5% CO_2_ for an additional 48 hours. Cells were harvested by centrifugation at 3,000 rpm, flash-frozen in liquid nitrogen, and stored at -80 °C until use.

Cell pellets were then lysed by Dounce homogenization in purification buffer (20 mM Tris, 150 mM NaCl and 5 mM DTT, pH 8.0) supplemented with aprotinin (2 µg/mL), leupeptin (1.4 µg/mL), and pepstatin A (0.5 µg/mL). Lysate was subjected to ultracentrifugation at 35,000 rpm for 40 minutes at 4 °C to isolate the membrane fraction. Membranes were solubilized for 2 hours at 4 °C in purification buffer supplemented with 1% w/v lauryl maltose neopentyl glycol (LMNG), 0.15% w/v cholesteryl hemisuccinate (CHS), and same concentration protease inhibitors. Following solubilization, the sample was ultracentrifuged again at 35,000 rpm for 40 minutes at 4 °C. The supernatant was collected and filtered through a 0.22 μm membrane before affinity purification. The filtered supernatant was loaded onto a streptavidin affinity column pre-equilibrated with purification buffer containing 0.1% w/v LMNG and 0.15% w/v CHS. Bound protein was eluted using the same buffer supplemented with 5 mM desthiobiotin. The eluate was concentrated by using a 30 kDa MWCO Amicon (Millipore) and further purified by size-exclusion chromatography using a Superose 6 Increase 10/300 GL column (GE Healthcare) in purification buffer containing 0.001% w/v LMNG and 0.0015% w/v CHS. Peak fractions were pooled for cryo-EM sample preparation. For VIAAT^PO4^, purification buffer was replaced with phosphate buffer (75 mM Na_2_HPO_4_, 25 mM NaH_2_PO_4_, pH 7.5). All other purification steps were identical.

### Cryo-EM sample preparation and data collection

Purified VIAAT protein was concentrated to approximately 10 mg ml⁻¹ using a 30 kDa MWCO Amicon (Millipore). For ligand-bound samples, the protein was incubated with 200 mM GABA or 200 mM glycine for 30 minutes on ice prior to grid preparation. Quantifoil R1.2/1.3 Cu 300-mesh grids (Quantifoil) were glow-discharged for 60 s under a hydrogen–oxygen atmosphere using a Solarus plasma cleaner (Gatan). A 2.5 μl aliquot of the protein sample was applied to the grids, incubated for 5 s, and blotted for 4–6 s at 4 °C under 100% humidity using a Vitrobot Mark IV (Thermo Fisher Scientific). The grids were then vitrified by plunge-freezing into liquid ethane cooled by liquid nitrogen. Data were collected on a Titan Krios transmission electron microscope (Thermo Fisher Scientific) operating at 300 kV, equipped with a K3 direct electron detector (Gatan) and a BioQuantum energy filter (Gatan) with a 20 eV slit width. Automated data acquisition was performed using EPU software (Thermo Fisher Scientific) at a nominal magnification of 105,000× in super-resolution mode, yielding a calibrated pixel size of 0.85 Å. Images were recorded with a defocus range of −1.2 to −2.2 μm. Each movie was dose-fractionated into 32 frames, with a total electron exposure of ∼60 e⁻ Å⁻².

### Cryo-EM data processing

For VIAAT^GABA^, VIAAT^Gly^, VIAAT^Apo^ and VIAAT^PO4^, we collected 3,017, 4,417, 2,238, 3,203 movies, respectively. Because the processing workflows were highly similar, the procedure for VIAAT^GABA^ is described as a representative example. All initial processing was performed in cryoSPARC^57^. Raw movie frames were motion-corrected, and contrast transfer function (CTF) parameters were estimated. Low-quality micrographs exhibiting contamination or poor CTF fits were manually removed. Particle picking was performed using the blob picker, yielding 3,191,330 particles from 2,936 micrographs, which were extracted with a box size of 256 pixels. Several rounds of 2D classification were carried out, and particles displaying well-defined features across a range of projection views were selected for ab initio reconstruction. Subsequent iterative 3D refinement, including heterogeneous refinement followed by non-uniform refinement, produced an initial 3.82 Å map from 97,694 particles. Particles corresponding to the best classes were then curated and used to train a Topaz neural network. Topaz-based picking, seeded with the initial 3D map, substantially enriched the particle dataset. Refinement of this enlarged particle pool yielded a 3.21 Å reconstruction from 319,903 particles. A mask generated from this map was applied during further 3D classification and local refinement, improving the resolution to 2.94 Å. Finally, particles were re-extracted with a 320-pixel box and subjected to mask-optimized local refinement, resulting in a final map at 2.7 Å resolution.

### Model building

The atomic model of the apo structure was built de novo using the high-resolution cryo-EM density map in COOT^58^. The map exhibited well-resolved features for the backbone and most side chains, enabling reliable model construction. Models of VIAAT^GABA^, VIAAT^GLy^, VIAAT^PO4^ were generated by rigid-body fitting of the apo model into the corresponding maps using UCSF Chimera^59^, followed by iterative manual rebuilding in COOT^58^ guided by the ligand densities and local conformational changes. Ligand restraint files were generated using the elBOW module in PHENIX^60^. All models were subjected to real-space refinement in PHENIX, incorporating secondary structure, rotamer, and geometry restraints. Model validation was performed using the cryo-EM validation module in PHENIX, evaluating global and local map-to-model correlation, MolProbity scores, clashscore, and Ramachandran statistics. All figures were prepared using open-source PyMOL^61^, UCSF Chimera^62^.

### Electrophysiological recording

Animal care and all experimental protocols were approved by the Animal Care and Use Committees at the Shenzhen Institute of Advanced Technology, Chinese Academy of Sciences (IACUC ID: SIAT-BSI-IRB-251017-NS-MKP-A0184). Primary hippocampal cultures were generated from newborn mice within 24 hours after birth. As previously described^63^, hippocampi were dissected out, and the dissociated cells were plated onto chemically stripped glass coverslips and maintained in a 37 °C tissue culture incubator. Cells were plated and maintained with tissue culture medium composed of Neurobasal-A (10888022) with 2% B-27 supplement (Thermo Fisher 17504044) and 1% GlutaMAX (Thermo Fisher 35050061).

To knock down endogenous VIAAT, a lentivirus expressing shRNA targeting the mRNA sequence 5’-GCTGGTGATGACGTGTATCTT-3’ was obtained from VectorBuilder (LV-U6-shRNA-CMV-GFP). The U6-shRNA-CMV-GFP cassette was subsequently replaced with either WT human VIAAT-IRES-mCherry or its mutant version to generate LV-hSyn-VIAAT (WT or mutant)-IRES-mCherry constructs. To protect the exogenous VIAAT from shRNA-mediated silencing, the sequence (5’-GCTGGTGATGACGTGCATCCTG-3’) in the human VIAAT complementary DNA (cDNA) was replaced with another (5’-GTTAGTTATGACTTGTATTTTA-3’) to produce a synonymous mutation for all VIAAT rescue constructs. All lentiviruses used to deliver rescue VIAAT were produced in-house by Ca^2+^-phosphate transfection in HEK293T cells. 48 h after transfection, the HEK293T cell supernatant was collected and stored frozen until further use. Both the shRNA and rescue lentiviruses were added to cultured neurons at DIV3.

Electrophysiological recordings in cultured hippocampal neurons were performed at DIV13-15. For IPSC recordings, the extracellular solution contained (in mM): 140 NaCl, 5 KCl, 2 CaCl_2_, 2 MgCl_2_, 10 glucose, 0.05 D-APV, 0.02 CNQX, 10 HEPES-NaOH (pH 7.4, ∼310 mOsm). Patch pipettes were pulled at 2-4 MΩ and filled with intracellular solution containing (in mM): 40 CsCl, 90 K-gluconate, 1.8 NaCl, 1.7 MgCl_2_, 3.5 KCl, 0.05 EGTA, 2 Mg-ATP, 0.4 Na_2_-GTP, 10 Phosphocreatine, 4 QX314-Cl, 10 HEPES-CsOH (pH 7.4, ∼300 mOsm). Cells were held at -70 mV during recording. All recordings were performed at room temperature (21-24 °C), and access resistance was monitored. Cells were discarded if the uncompensated access exceeded 15 MΩ, and access resistance was compensated to 3-4 MΩ during recording. A custom-made bipolar focal stimulation electrode made from nichrome wire was used to evoke action potentials. Data acquisition was performed with an Axon Multiclamp 700B amplifier and digitized with a Digidata 1440A digitizer. All data analyses were done with pClamp10. Quantification of peak amplitudes was performed by subtracting the baseline current and finding the negative peak of the response following the stimulus artifact.

### Quantification of VIAAT expression level

To quantify the expression level of VIAAT delivered via lentivirus, confocal imaging was performed following standard protocols. Cultured neurons grown on glass coverslips were washed with PBS, fixed with 4% PFA for 10 minutes, rinsed again with PBS, and then blocked and permeabilized in 0.05% Triton X-100 with 3% BSA in PBS (TBP) for 1 hour. Primary antibodies were diluted in TBP, and coverslips were incubated overnight at 4 °C. The primary antibodies used included rabbit anti-VIAAT (1:500, Synaptic Systems, #131003) and guinea pig anti-Synaptophysin (1:500, Synaptic Systems, #101004). After primary antibody incubation, coverslips were washed three times for 5 minutes each in TBP.

Secondary antibodies conjugated to Alexa Fluor 555 and 633 were applied at a 1:500 dilution in TBP and incubated for 1 hour at room temperature. Following secondary staining, coverslips were washed three times for 5 minutes each in TBP, rinsed once with deionized water, air-dried, and mounted onto glass slides.

Images were acquired using a Leica Stellaris 5 SR confocal microscope equipped with a 63× oil immersion objective (1.44 numerical aperture). For quantification of synaptic VIAAT expression, regions of interest (ROIs) were defined based on Synaptophysin puncta using the “Analyze Particles” function in ImageJ. The average fluorescence intensities of VIAAT and Synaptophysin within these ROIs were measured for each image (Extended Data Fig. 8b). The VIAAT intensity was then normalized to the Synaptophysin intensity within the same ROIs (Extended Data Fig. 8c). The IPSC amplitudes recorded from neurons expressing wild-type or mutant VIAAT were normalized to their respective expression levels.

### Molecular dynamics simulations

To delineate the roles of ions and residue protonation in modulating GABA binding and release, we conducted a series of molecular dynamics simulations across defined ionic conditions while systematically varying the protonation states of key residues. The cryo-EM structure of the VIAAT in complex with GABA served as the initial structure for molecular dynamics simulations. Each VIAAT^GABA^ complex was embedded into a lipid bilayer consisting of 128 1-palmitoyl-2-oleoyl-sn-glycero-3-phosphocholine (POPC) molecules and 32 cholesterol molecules via the CHARMM-GUI web server^64–66^. The assembled systems were solvated in ∼18,000 TIP3P water molecules, neutralized, and adjusted to a final ionic strength of 150 mM NaCl. The final simulation boxes measured roughly 8.24 × 8.24 × 12 nm³. All molecular dynamics simulations were performed using GROMACS v.2024.3 with the CHARMM36 force field^67^ for protein and lipids and CGenFF^68^ for GABA. Periodic boundary conditions (PBC) were applied throughout all simulations. Lennard-Jones (LJ) interactions were truncated at 10 Å and smoothed to zero between 10 and 12 Å. Electrostatic interactions were calculated using the Particle Mesh Ewald (PME) method, with a grid spacing of 1.2 Å. The H-bond was constrained using the LINCS algorithm^69^. A 2 fs time step was utilized with temperature maintained at 310 K via the Nose−Hoover algorithm^70^, and the pressure was controlled semi-isotropically at 1 atm using the Parrinello−Rahman barostat^71^. Prior to production simulations, each system was energy-minimized using the steepest descent algorithm, followed by sequential 10 ns NVT and NPT equilibration phases to relax the solvent and lipid bilayer. Production molecular dynamics simulations were then performed for 400 ns, with trajectory frames saved every 1ns for downstream analysis. Clustering analysis was carried out using the GROMOS software^72^.

## Supporting information

This document contains Extended Data Figure 1-12 and Extended Data Table 1-2.

## Data availability

The three-dimensional cryo-EM density maps of human VIAAT in the apo state (NaCl buffer), GABA-bound state (NaCl buffer), glycine-bound state (NaCl buffer), and apo state prepared in phosphate buffer have been deposited in the Electron Microscopy Data Bank (https://www.ebi.ac.uk/emdb/) under the accession codes EMD-67826, EMD-67799, EMD- 67798, and EMD-67797, respectively. The corresponding atomic coordinates for VIAAT in these states have been deposited in the Protein Data Bank (https://www.rcsb.org/) under the accession codes 21MN, 21LK, 21LJ, and 21LI, respectively.

## Author contribution

Y.Z. conceived and supervised the project. T.H. and K.H. prepared cryo-EM samples. K.M. performed functional assays. Z.C. and Q.B. conducted MD simulations. J.Z., R.L., H.Z., and Y.L. carried out cryo-EM data collection. K.H., T.H., Y.W., and Q.C. processed cryo-EM data and built models. Y.W., K.H., T.H., and X.X. analyzed the structures and prepared the figures. Y.Z., T.H., K.H., Y.W., K.M., and H.Z. wrote and revised the manuscript.

## Acknowledgments

We thank B. Xu at Peking University Institute of Advanced Agricultural Sciences for support in cryo-EM data collection. This work is funded by Young Scientists Fund (A) of the National Natural Science Foundation of China (32525036 to Y.Z.), Brain Science and Brain-like Intelligence Technology - National Science and Technology Major Project (2022ZD0205800 to Y.Z.), Chinese Academy of Sciences Project for Young Scientists in Basic Research (YSBR-104), National Key R&D Program of China (2021YFA1301501 to Y.Z.), National Natural Science Foundation of China (92157102 to Y.Z.), Beijing Anding hospital, Capital Medical University (YG202503 to Y.Z.), Shenzhen Medical Research Fund (A2503041 to K.M.), High-level Talent Research Start-up Fund of China Pharmaceutical University (3150010178 to T.H.), China National Postdoctoral Program for Innovative Talents (BX2026120 to Y.W.), and Key Laboratory of Anesthesiology and Resuscitation (Huazhong University of Science and Technology), Ministry of Education (2025MZFS001).

