## Supplementary material for "Substrate recognition and ion coupling mechanism of the human VIAAT": This document contains Extended Data Figure 1-12 and Extended Data Table 1-2.

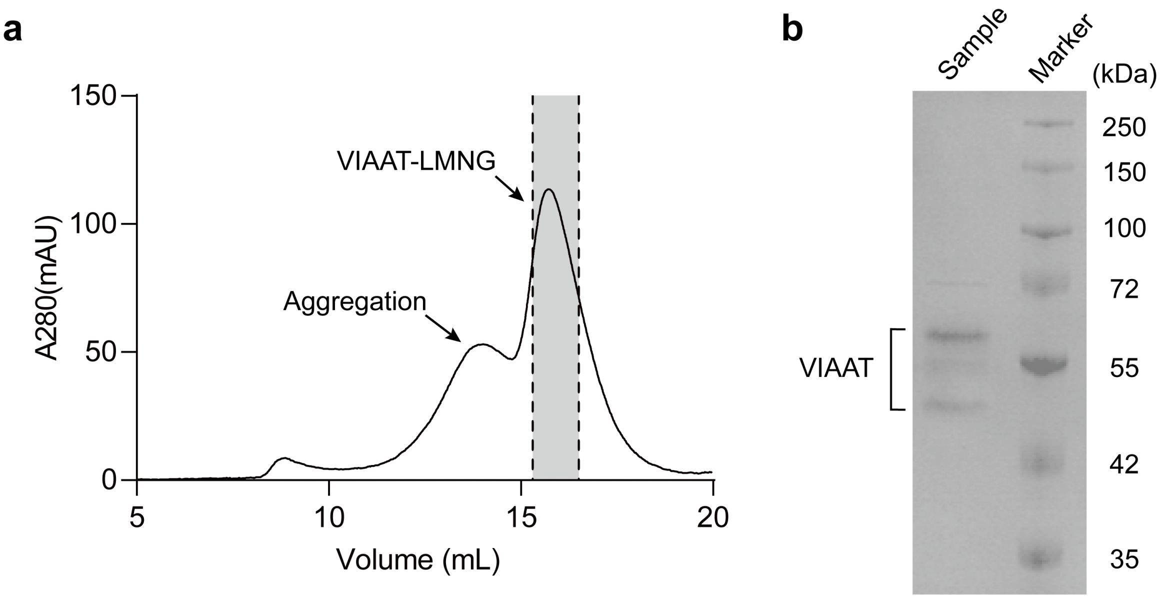

Extended Data Figure 1. Purification and biochemical characterization of VIAAT.

**a,** Size-exclusion chromatography (SEC) profile of VIAAT purified in LMNG. Absorbance at 280 nm (A280) is plotted as a function of elution volume (mL). The main peak corresponding to monodisperse VIAAT–LMNG is indicated, while the shoulder peak represents aggregated species. Peak fractions (shaded) were pooled and concentrated for subsequent cryo-EM sample preparation.

**b,** Coomassie blue–stained SDS–PAGE analysis of the VIAAT cryo-EM sample. Molecular weight markers (kDa) are shown on the right, and the VIAAT band is indicated.

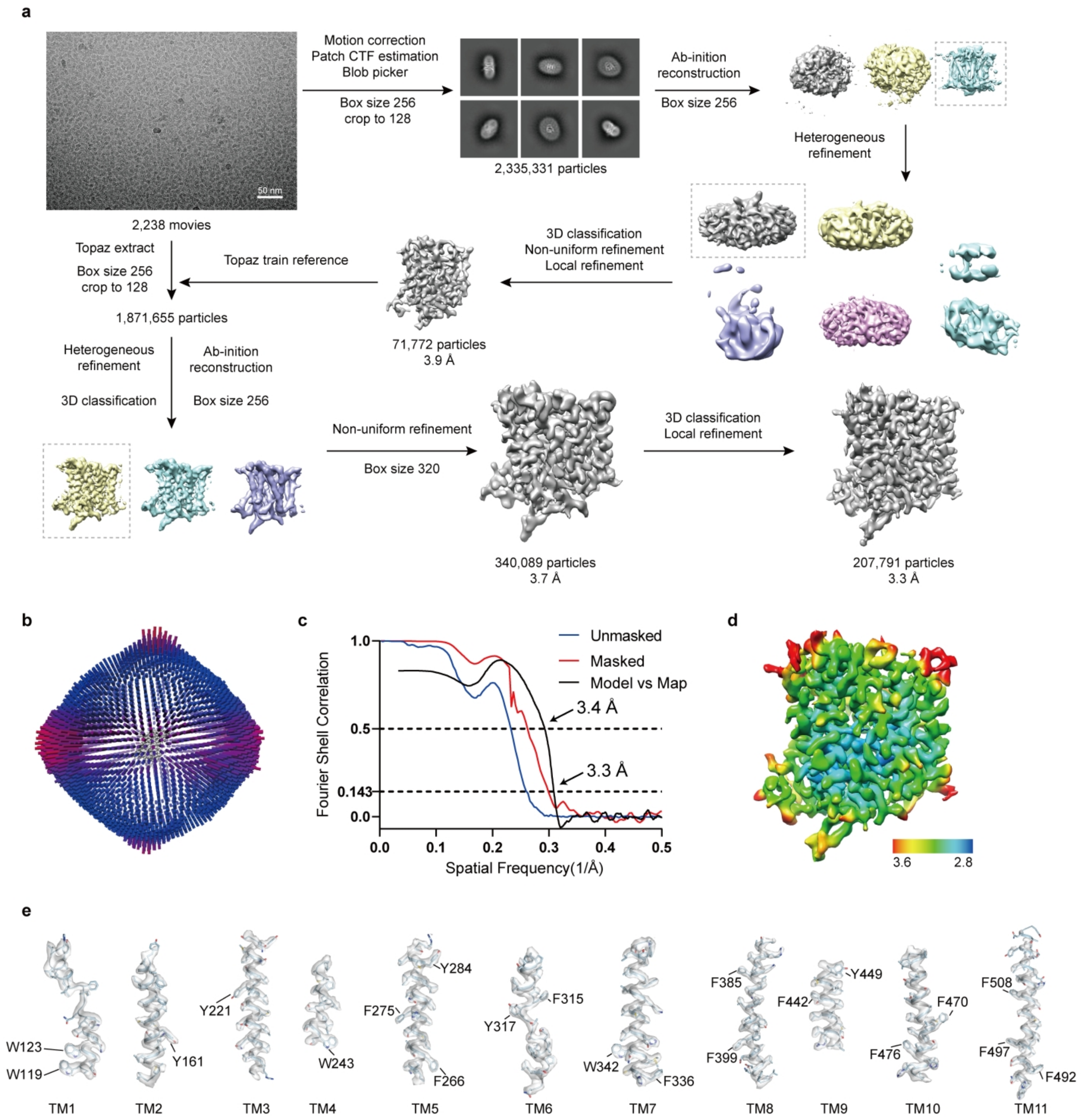

### Extended Data Figure 2. Cryo-EM data processing of VIAAT$^{Apo}$.

**a**, Flowchart for cryo-EM data processing of VIAAT$^{Apo}$. **b**, Angular distribution of particles contributing to the final reconstruction. **c**, Fourier shell correlation (FSC) curves of the unmasked (blue) and masked (red) half-maps, plotted as a function of spatial frequency (1/Å). The final resolution is estimated at 3.3 Å based on the gold-standard FSC criterion. **d**, Final cryo-EM density map colored by local resolution, ranging from 2.8 Å (blue) to 3.6 Å (red). **e**, Representative cryo-EM density with the fitted atomic model for transmembrane helices TM1–TM11 of VIAAT$^{Apo}$.

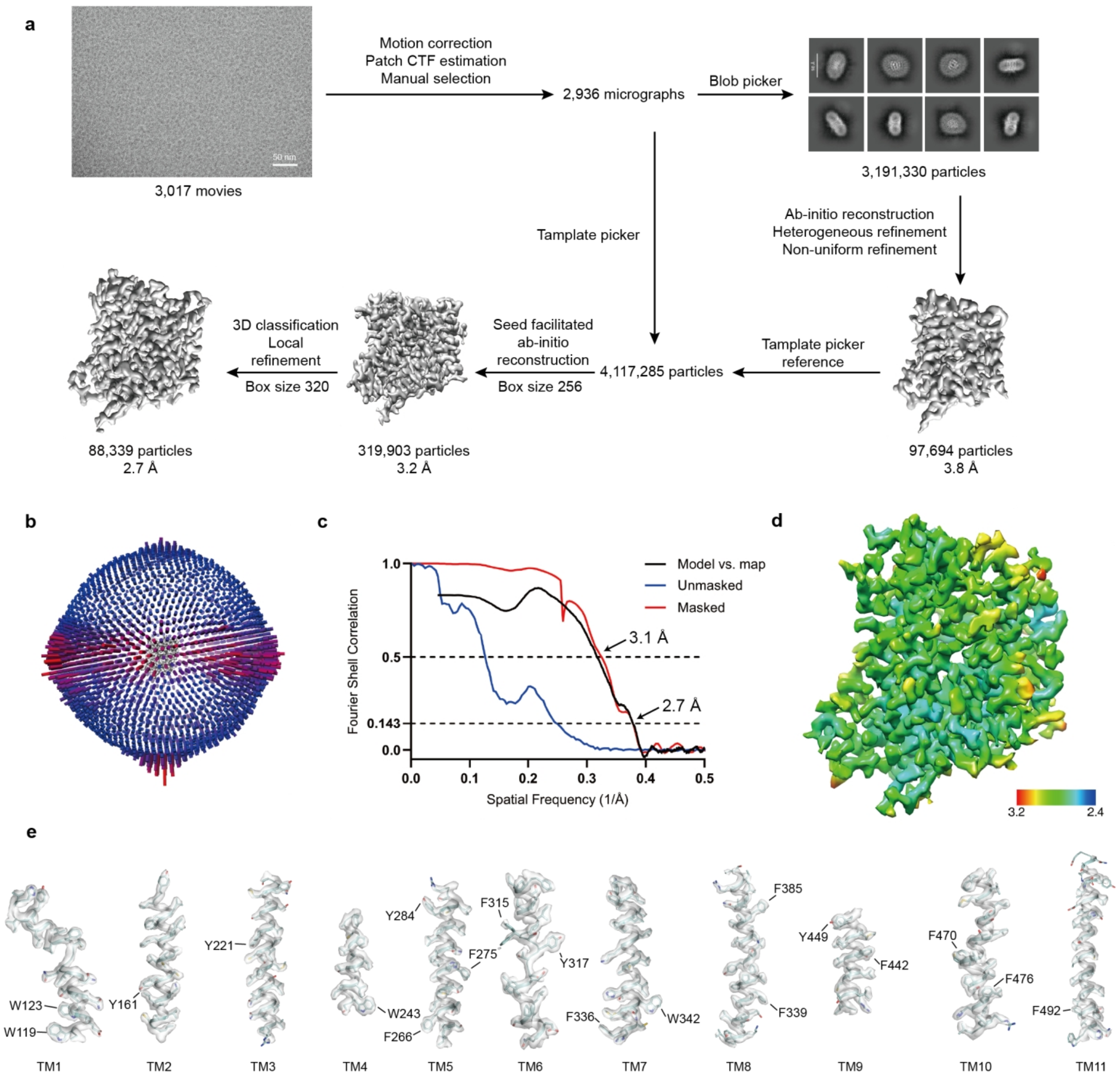

### Extended Data Figure 3. Cryo-EM data processing of VIAAT$^{GABA}$.

**a**, Flowchart for cryo-EM data processing of VIAAT$^{GABA}$. **b**, Angular distribution of the particles used in the final reconstruction. **c**, Fourier shell correlation (FSC) curves of the unmasked (blue) and masked (red) half-maps, plotted as a function of spatial frequency (1/Å). The final resolution is estimated at 2.7 Å based on the gold-standard FSC criterion. **d**, Final cryo-EM density map colored by local resolution, ranging from 2.4 Å (blue) to 3.2 Å (red). **e**, Representative cryo-EM density with the fitted atomic model for transmembrane helices TM1–TM11 of VIAAT$^{GABA}$.

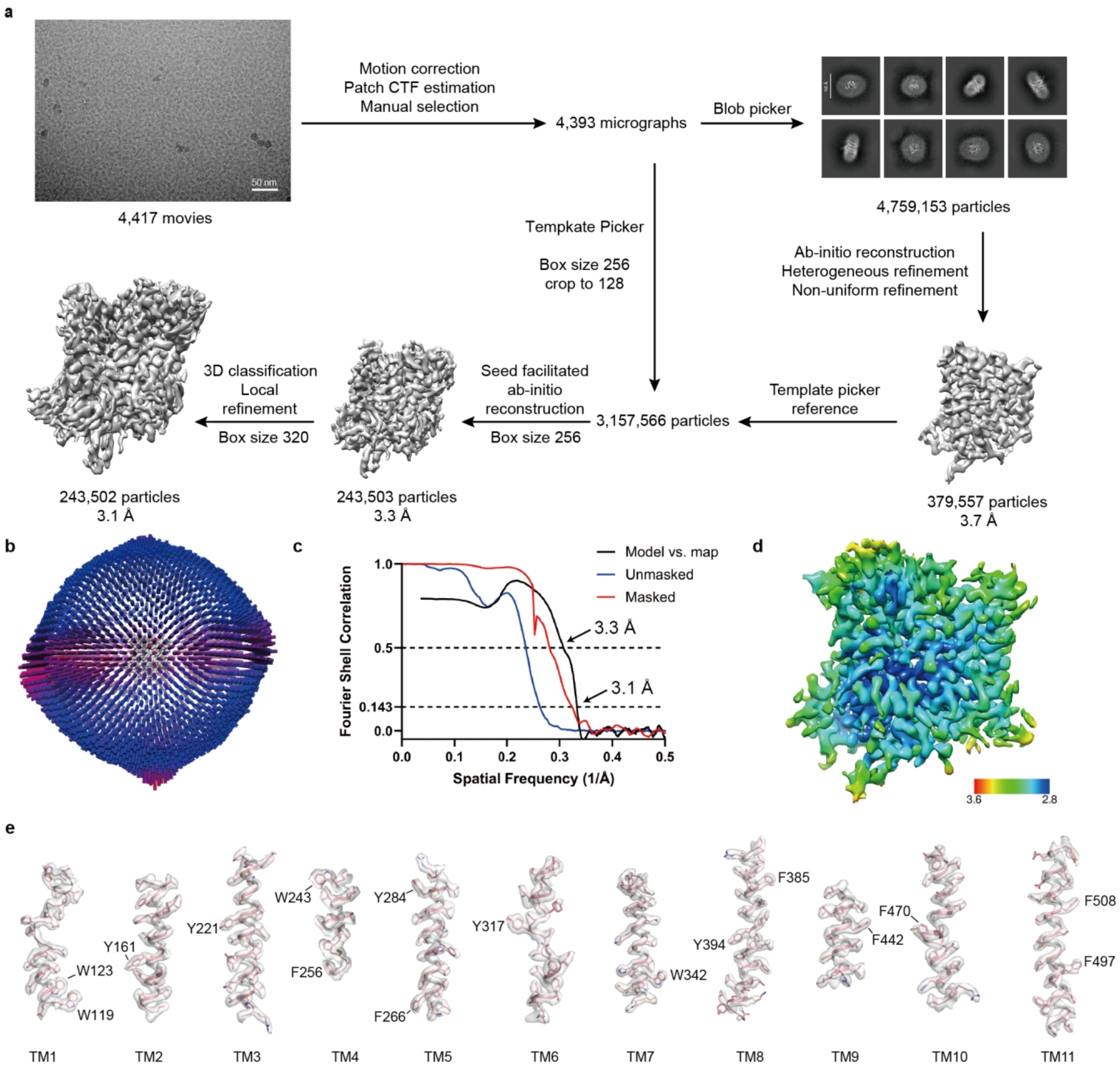

### Extended Data Figure 4. Cryo-EM data processing of VIAAT$^{Gly}$.

**a**, Flowchart for cryo-EM data processing of VIAAT$^{Gly}$. **b**, Angular distribution of the particles used in the final reconstruction. **c**, Fourier shell correlation (FSC) curves of the unmasked (blue) and masked (red) half-maps, plotted as a function of spatial frequency (1/Å). The final resolution is estimated at 3.1 Å based on the gold-standard FSC criterion. **d**, Final cryo-EM maps colored by local resolution, scale ranging from 2.8 Å (blue) to 3.6 Å (red). **e**, The cryo-EM density and atomic model of TM1–TM11 of VIAAT$^{Gly}$.

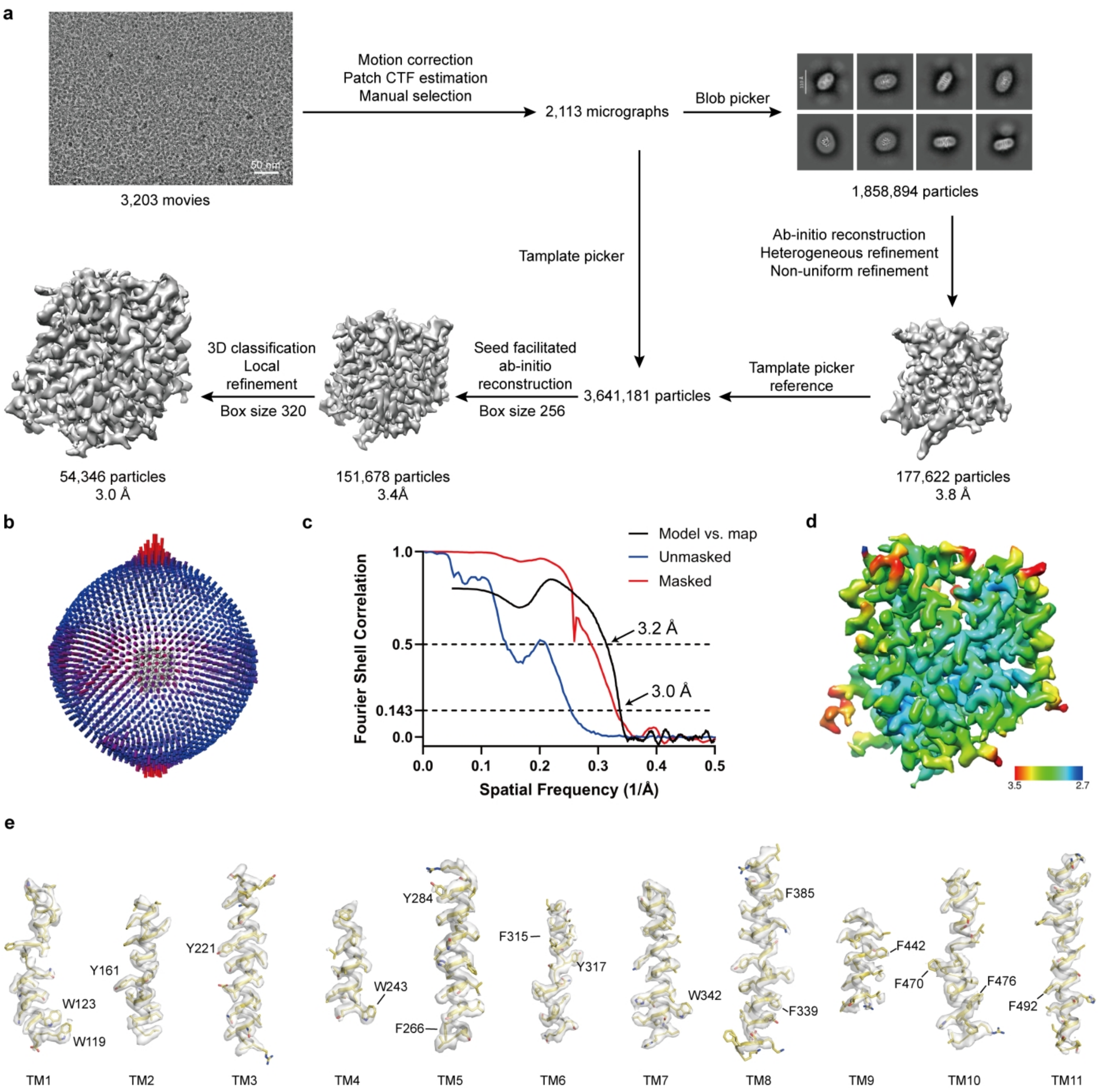

### Extended Data Figure 5. Cryo-EM data processing of VIAAT$^{PO4}$.

**a**, Flowchart for cryo-EM data processing of VIAAT$^{PO4}$. **b**, Angular distribution of the particles used in the final reconstruction. **c**, Fourier shell correlation (FSC) curves of the unmasked (blue) and masked (red) half-maps, plotted as a function of spatial frequency (1/Å). The final resolution is estimated at 3.0 Å based on the gold-standard FSC criterion. **d**, Final cryo-EM density map colored by local resolution, ranging from 2.7 Å (blue) to 3.5 Å (red). **e**, Representative cryo-EM density with the fitted atomic model for transmembrane helices TM1–TM11 of VIAAT$^{PO4}$.

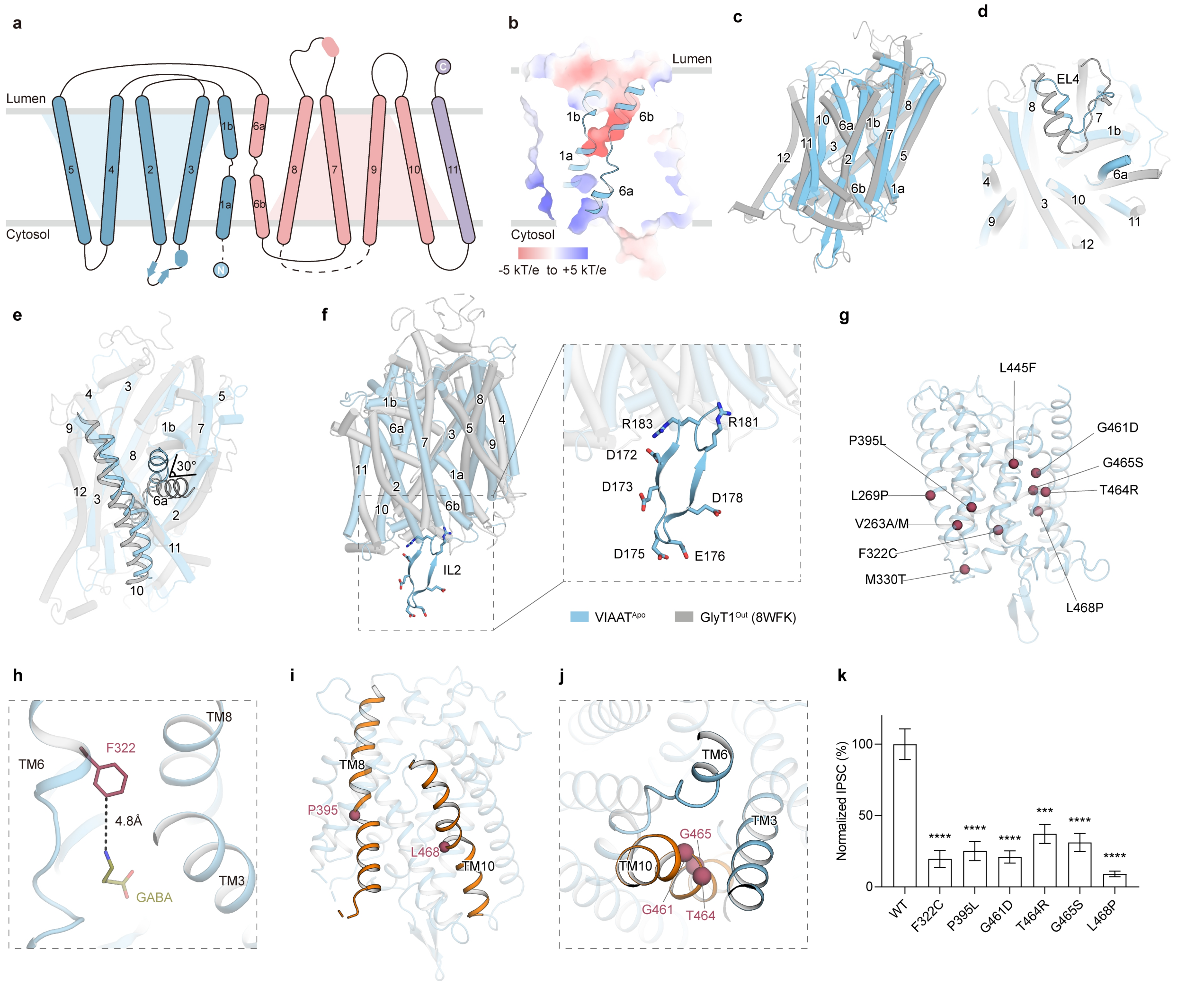

### Extended Data Figure 6. Structural organization of VIAAT and mapping of disease-associated mutations.

**a,** Topology of VIAAT. TM1–TM5, TM6–TM10, and TM11 are colored light blue, light pink, and lavender, respectively. Loop regions are shown as lines, whereas unresolved regions are indicated by dashed lines. The luminal and cytosolic sides are labeled.

**b,** Electrostatic surface representation of VIAAT$^{Apo}$. TM1 and TM6 are shown as blue cartoons. Positively and negatively charged surfaces are colored blue and red, respectively.

**c**, Side view of the structural alignment between VIAAT$^{Apo}$ (cyan) and GlyT1$^{Out}$ (PDB: 8WFK; grey), superposed based on the scaffold domain. Overall structures are shown in cartoon representation, with transmembrane helices labeled.

**d,e**, Top-down view (d) and side view (e) of the superposed VIAAT$^{Apo}$ and GlyT1$^{Out}$ structures highlighting conformational differences. Regions exhibiting pronounced structural divergence are shown without transparency.

**f,** Comparison of the loop between TM2 and TM3 in VIAAT$^{Apo}$ (cyan) and GlyT1$^{Out}$ (grey). This loop is substantially larger in VIAAT and is enriched in charged residues compared with the corresponding region in GlyT1.

**g,** Mapping of epilepsy-associated missense variants onto the $VIAAT^{Apo}$ structure. Cα atoms of pathogenic mutations are shown as red spheres.

**h**, Close-up view of the pathogenic mutation F322C. Residue F322 and bound GABA are shown as raspberry and olive sticks, respectively, with the distance between F322 and GABA indicated.

**i**, Locations of the pathogenic mutations P395L and L468P. TM8 and TM10 are colored orange, while other helices are shown in cyan. P395 and L468 are depicted as raspberry spheres.

**j,** Close-up view of pathogenic mutation sites on TM10. Cα atoms of G461, T464, and G465 are shown as raspberry spheres.

**k**, IPSC amplitudes of disease-associated VIAAT variants were compared with WT VIAAT values, and then corrected for the corresponding synaptic expression levels. Data are shown as the mean ± SEM from biologically independent experiments ($n$ = 8–11). Statistical significance was assessed by unpaired $t$-test with $\alpha = 0.05$. *$P < 0.05$; **$P < 0.01$; ***$P < 0.001$; ****$P < 0.0001$ versus WT VIAAT.

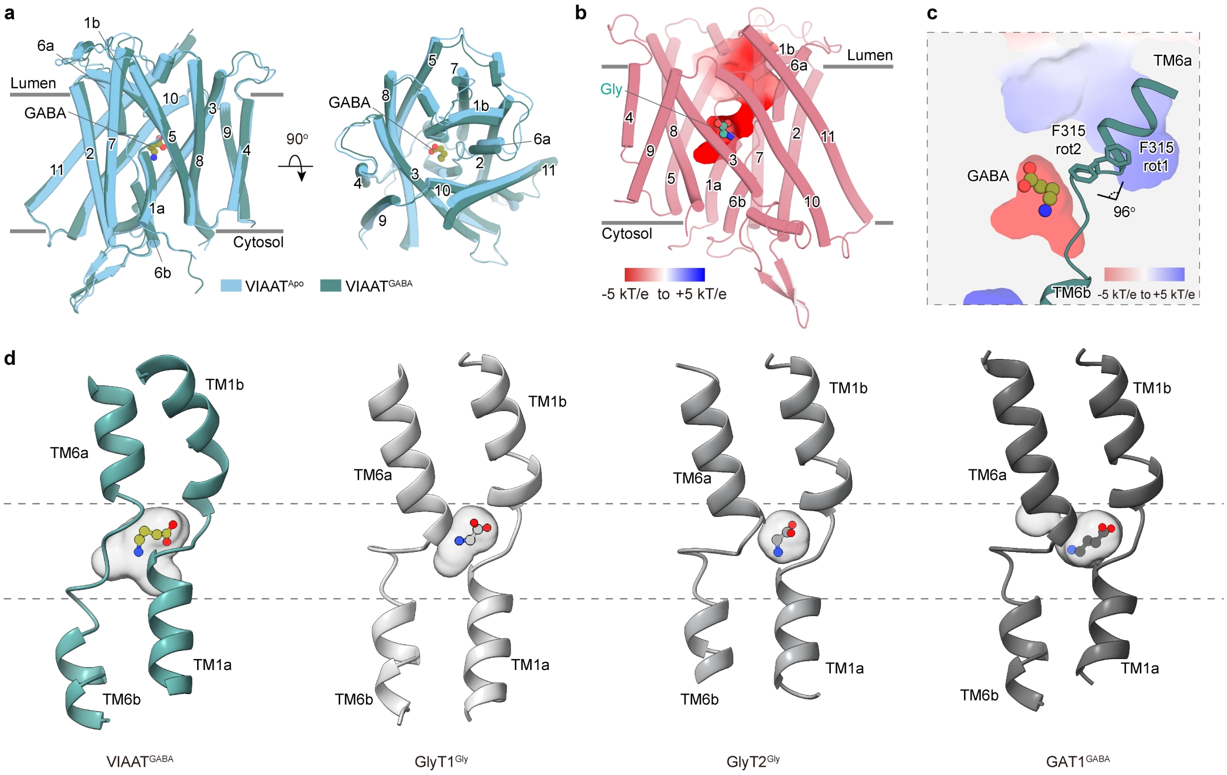

### Extended Data Figure 7. Comparison of the substrate-binding pocket in VIAAT with neurotransmitter sodium symporters.

**a**, Overall structures of VIAAT$^{Apo}$ and VIAAT$^{GABA}$, superposed using the scaffold domain (TM3, TM4, TM8, and TM9), are shown as cylinders.

**b**, Overall structure of VIAAT$^{Gly}$ highlighting the luminal entrance and the glycine-binding site. Electrostatic surface potential is shown, with positively and negatively charged regions colored blue and red, respectively.

**c**, Close-up view of the F315 conformation in the VIAAT$^{GABA}$ structure. Residue F315 is shown as sticks and GABA as spheres. Nitrogen and oxygen atoms are colored blue and red, respectively. The angle between the two alternative F315 rotamers is indicated. Electrostatic surface potential is shown, with positively and negatively charged regions colored blue and red, respectively.

**d**, Comparison of the central substrate-binding pocket of VIAAT with those of GlyT1, GlyT2, and GAT1. The binding pockets are shown as grey surfaces. Substrate-binding pockets of these transporters are visualized using PyVOL with a minimum radius of 1.8 Å and a maximum radius of 3.4 Å.

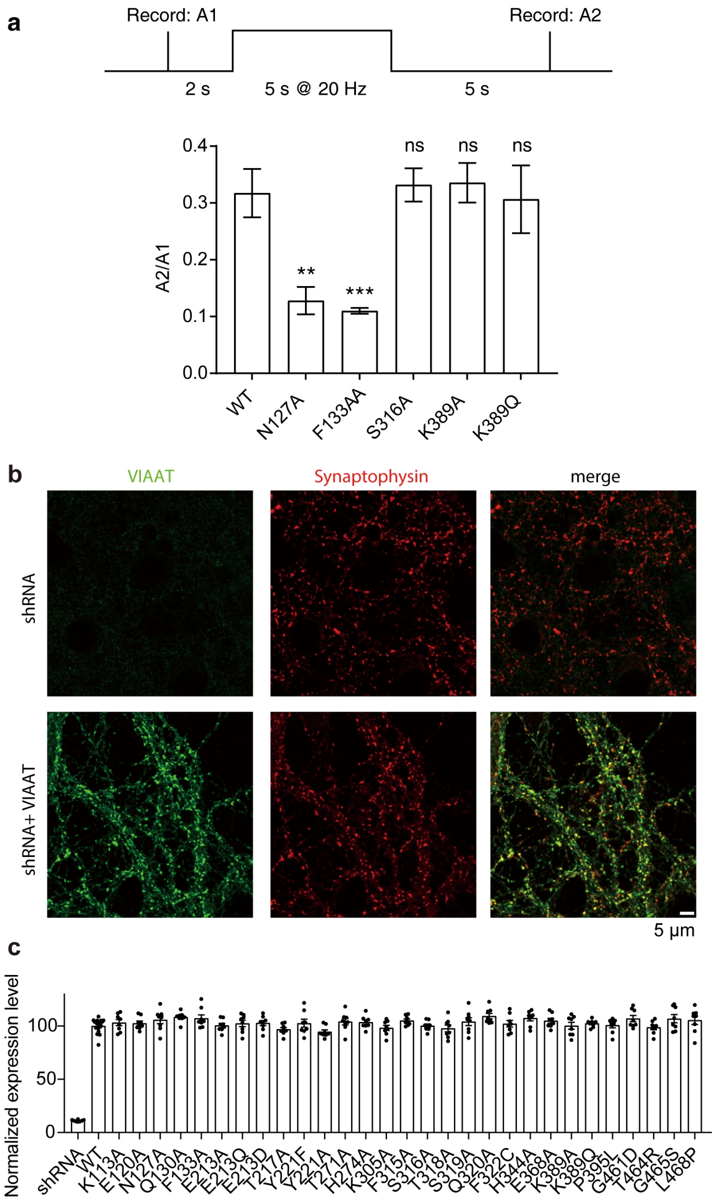

### Extended Data Figure 8. IPSC recovery kinetics following train stimulation and synaptic expression analysis of wild-type and mutant VIAAT.

**a**, Recovery of IPSCs evoked by VIAAT after train stimulation. Data are shown as the mean ± SEM from biologically independent experiments ($n$ = 7–12). Statistical significance was assessed using an unpaired $t$-test with α = 0.05. *$P$ < 0.05; **$P$ < 0.01; ***$P$ < 0.001; ****$P$ < 0.0001. **b**, DIV13-15 mouse primary hippocampal neurons were transfected with lentivirus containing $shRNA^{VIAAT}$ alone or co-transduced with a VIAAT rescue construct and immunostained for Synaptophysin. Representative confocal microscopy images are shown. **c**, Synaptic VIAAT expression was quantified within Synaptophysin-defined ROIs, and VIAAT fluorescence was normalized to Synaptophysin. The resulting expression levels were used to normalize IPSC amplitudes of WT and mutant VIAAT.

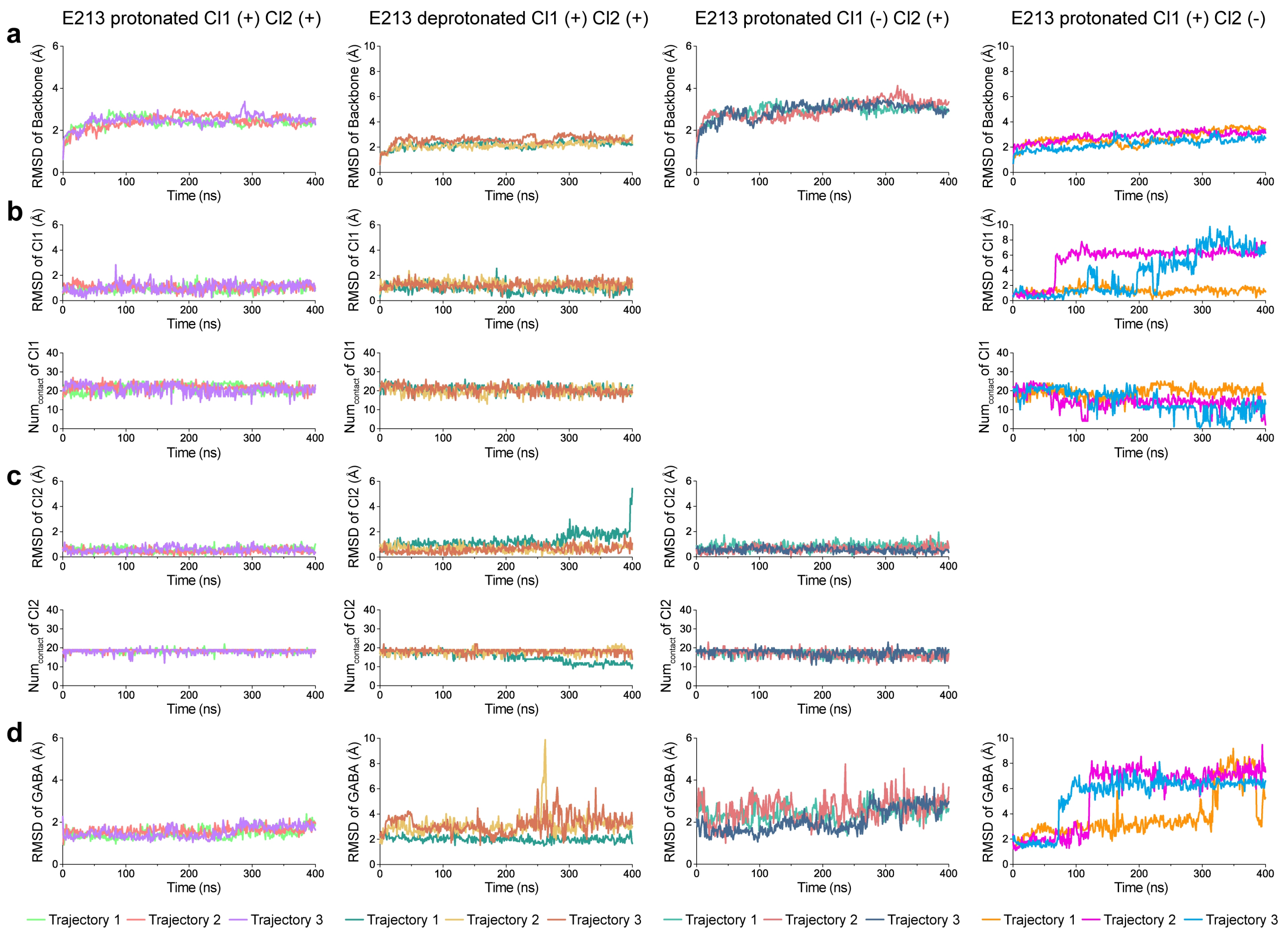

### Extended Data Figure 9. Effects of different conditions on GABA and ion stability.

**a**, The root-mean-square deviation (RMSD) of the VIAAT protein backbone Cα atoms. For all RMSD calculations, the initial structure was used as the reference. Results from the three independent trajectories are distinguished by distinct colors.

**b**, (Top) The RMSD of Cl1 ion. (Bottom) The atom number of contacts ($Num_{contact}$) between the Cl1 ion and its coordinating residues (T271, H274, K389).

**c**, (Top) The RMSD of Cl2 ion. (Bottom) The atom number of contacts ($Num_{contact}$) between the Cl2 ion and its coordinating residues (S319, Q320, H344).

**d**, The RMSD of GABA heavy atoms.

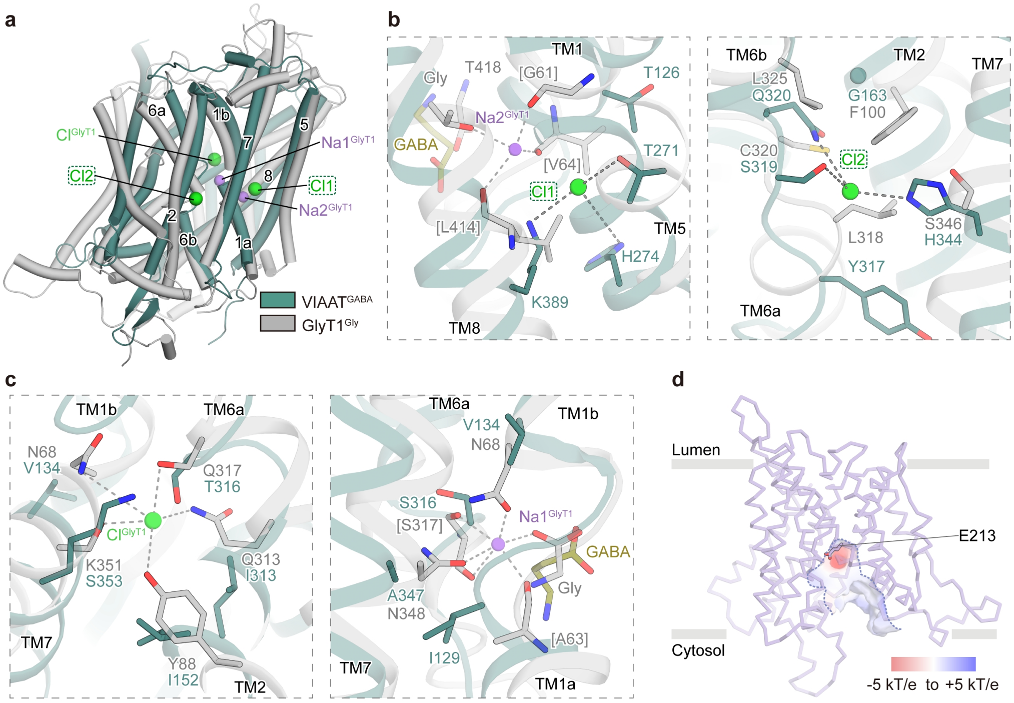

### Extended Data Figure 10. Structural comparison of ion-binding sites between VIAAT and GlyT1.

**a,** Structural superposition of VIAAT$^{GABA}$ (dark green) and GlyT1$^{Gly}$ (grey), aligned based on the scaffold domain. Chloride ions and chloride-binding sites are shown as green spheres, and sodium ions are shown as purple spheres.

**b,** Comparison of the Cl1 and Cl2 binding sites in VIAAT$^{GABA}$ and corresponding sites in GlyT1$^{Gly}$. GABA is shown as yellow sticks. Chloride ions and sodium ions are depicted as green and purple spheres, respectively. Key coordinating residues are shown as sticks.

**c,** Structural comparison of the chloride-binding site (left) and Na1-binding site (right) in GlyT1 (grey) and corresponding position in VIAAT (dark green). The residues coordinated ion binding in glyT1 and corresponding residues in VIAAT are shown as sticks. Chloride and sodium ion are displayed as green and purple spheres, respectively.

**d,** Position of E213 in the cytosol-facing homology model of VIAAT. The cytosol-exposed pocket is outlined with dashed lines, and the electrostatically negative region is shown in red.

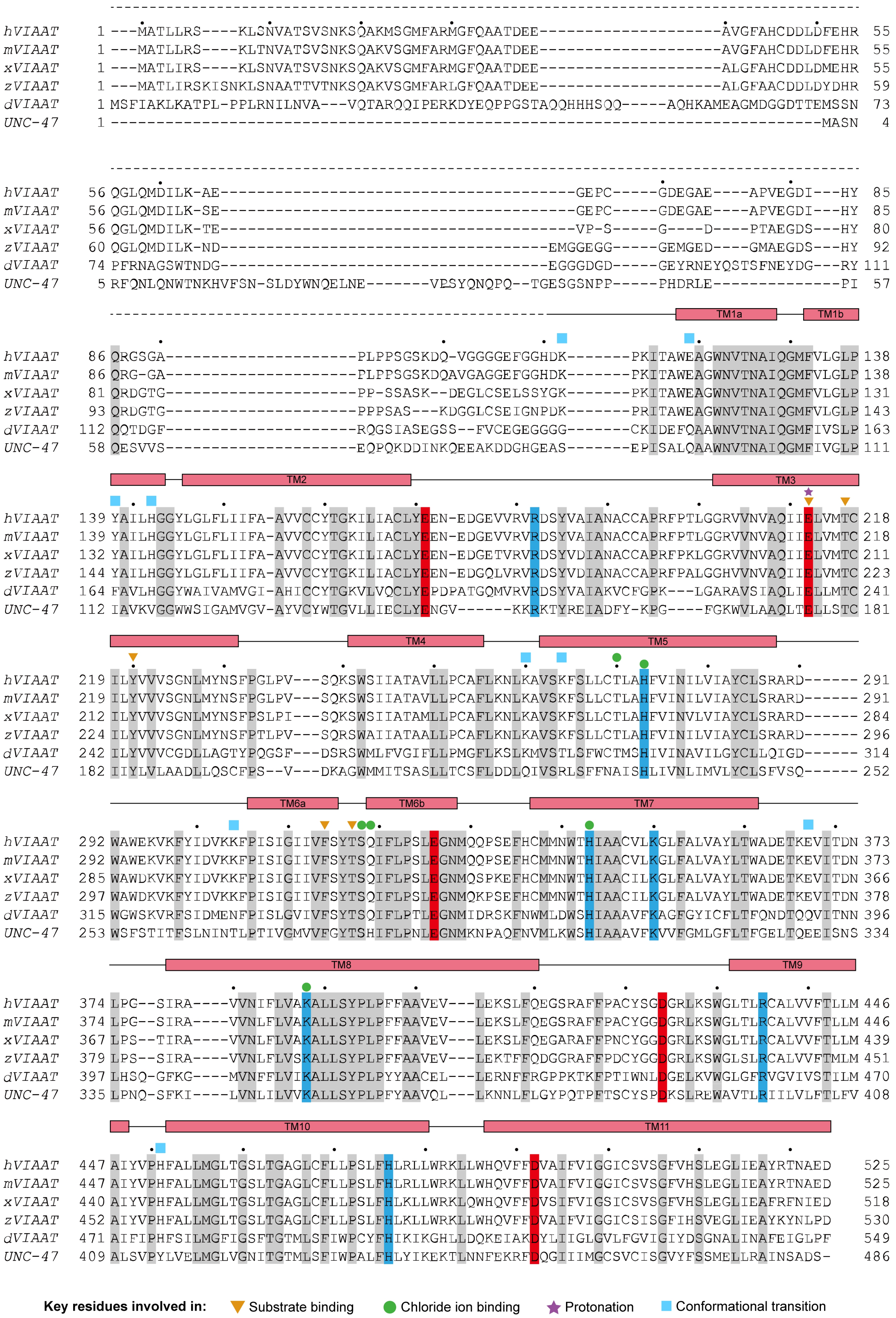
hVIAAT 1 ---MATLLRS----KLSNVATSVSNKSQAKMSGMFARMGFQAATDEE--------------------AVGFAHCDDLDFEHR 55
mVIAAT 1 ---MATLLRS----KLTNVATSVSNKSQAKVSGMFARMGFQAATDEE--------------------AVGFAHCDDLDFEHR 55
xVIAAT 1 ---MATLIRS----KLSNVATSVSNKSQAKVSGMFARMGFQAATDEE--------------------ALGFAHCDDLDMEHR 55
zVIAAT 1 ---MATLIRSKISNKLSNAATTVTNKSQAKVSGMFARLGFQAATDEE--------------------ALGFAACDDLDYDHR 59
dVIAAT 1 MSFIAKLKATPL-PPLRNILNVA---VQTARQQIPERKDYEQPPGSTAQQHHHSQQ-----AQHKAMEAGMDGGDTTEMSSN 73
UNC-47 1 -------------------------------------------------------------------------------MASN 4
hVIAAT 56 QGLQMDILK-AE---------------------------------------GEPC-----GDEGAE----APVEGDI---HY 85
mVIAAT 56 QGLQMDILK-SE---------------------------------------GEPC-----GDEGAE----APVEGDI---HY 85
xVIAAT 56 QGLQMDILK-TE---------------------------------------VP-S-----G----D----PTAEGDS---HY 80
zVIAAT 60 QGLQMDILK-ND-------------------------------------EMGGEGG-----GEMGED----GMAEGDS---HY 92
dVIAAT 74 PFRNAGSWTNDG-------------------------------------EGGGDGD-----GEYRNEYQSTSFNEYDG---RY 111
UNC-47 5 RFQNLQNWTNKHVFSN-SLDYWNQELNE-------VESYQNQPQ--TGESGSNPP-----PHDRLE--------------PI 57
TM1a
TM1b
hVIAAT 86 QRGSGA---------------------PLPPSGSKDQ-VGGGGEFGGHDK-------PKITAWEAGWNVTNAIQGMFVLGLP 138
mVIAAT 86 QRG-GA---------------------PLPPSGSKDQAVGAGGEFGGHDK-------PKITAWEAGWNVTNAIQGMFVLGLP 138
xVIAAT 81 QRDGTG---------------------PP-SSASK--DEGLCSELSSYGK-------PKITAWEAGWNVTNAIQGMFVLGLP 131
zVIAAT 93 QRDGTG---------------------PPPSAS---KDGGLCSEIGNPDK-------PRITAWEAGWNVTNAIQGMFVLGLP 143
dVIAAT 112 QQTDGF---------------------RQGSIASEGSS--FVCEGEGGGG-------CKIDEFQAAWNVTNAIQGMFIVSLP 163
UNC-47 58 QESVVS---------------------EQPQKDDINKQEEAKDDGHGEAS-------EPISALQAAWNVTNAIQGMFIVGLP 111
TM2
TM3
hVIAAT 139 YAILHGGYLGLFLIIFA-AVVCCYTGKILIACLYEEN-EDGEVVRVRDSYVAIANACCAPRFPTLGGRVVNVAQIIELVMTC 218
mVIAAT 139 YAILHGGYLGLFLIIFA-AVVCCYTGKILIACLYEEN-EDGEVVRVRDSYVAIANACCAPRFPTLGGRVVNVAQIIELVMTC 218
xVIAAT 132 YAILHGGYLGLFLIIFA-AVVCCYTGKILIACLYEEN-EDGETVRVRDSYVDIANACCAPRFPKLGGRVVNVAQIIELVMTC 211
zVIAAT 144 YAILHGGYLGLFLIIFA-AVVCCYTGKILIACLYEEN-EDGQLVRVRDSYVDIANACCAPRFPALGGHVVNVAQIIELVMTC 223
dVIAAT 164 FAVLHGGYWAIVAMVGI-AHICCYTGKVLVQCLYEPDPATGQMVRVRDSYVAIAKVCFGPK---LGARAVSIAQLIELLMTC 241
UNC-47 112 IAVKVGGWWSIGAMVGV-AYVCYWTGVLLIECLYENGV------KKRKTYREIADFY-KPG----FGKWVLAAQLTELLSTC 181
TM4
TM5
hVIAAT 219 ILYVVVSGNLMYNSFPGLPV---SQKSWSIIATAVLLPCAFLKNLKAVSKFSLLCTLAHFVINILVIAYCLSRARD------ 291
mVIAAT 219 ILYVVVSGNLMYNSFPGLPV---SQKSWSIIATAVLLPCAFLKNLKAVSKFSLLCTLAHFVINILVIAYCLSRARD------ 291
xVIAAT 212 ILYVVVSGNLMYNSFPSLPI---SQKSWSIIATAMLLPCAFLKNLKAVSKFSLLCTLAHFVINVLVIAYCLSRARD------ 284
zVIAAT 224 ILYVVVSGNLMYNSFPTLPV---SQRSWAIIATAALLPCAFLKNLKAVSKFSLLCTLAHFVINILVIAYCLSRARD------ 296
dVIAAT 242 ILYVVVCGDLLAGTYPQGSF---DSRSWMLFVGIFLLPMGFLKSLKMVSTLSFWCTMSHIVINAVILGYCLLQIGD------ 314
UNC-47 182 IIYLVLAADLLQSCFPS--V---DKAGWMMITSASLLTCSFLDDLQIVSRLSFFNAISHLIVNLIMVLYCLSFVSQ------ 252
TM6a
TM6b
TM7
hVIAAT 292 WAWEKVKFYIDVKKFPISIGIIVFSYTSQIFLPSLEGNMQQPSEFHCMMNWTHIAACVLKGLFALVAYLTWADETKEVITDN 373
mVIAAT 292 WAWEKVKFYIDVKKFPISIGIIVFSYTSQIFLPSLEGNMQQPSEFHCMMNWTHIAACVLKGLFALVAYLTWADETKEVITDN 373
xVIAAT 285 WAWDKVKFYIDVKKFPISIGIIVFSYTSQIFLPSLEGNMQSPKEFHCMMNWTHIAACILKGLFALVAYLTWADETKEVITDN 366
zVIAAT 297 WAWDKVKFYIDVKKFPISIGIIVFSYTSQIFLPSLEGNMQKPSEFHCMMNWTHIAACILKGLFALVAYLTWADETKEVITDN 378
dVIAAT 315 WGWSKVRFSIDMENFPISLGVIVFSYTSQIFLPTLEGNMIDRSKFNWMLDWSHIAAAVFKAGFGYICFLTFQNDTQQVITNN 396
UNC-47 253 WSFSTITFSLNINTLPTIVGMVVFGYTSHIFLPNLEGNMKNPAQFNVMLKWSHIAAAVFKVVFGMLGFLTFGELTQEEISNS 334
TM8
TM9
hVIAAT 374 LPG--SIRA----VVNIFLVAKALLSYPLPFFAAVEV---LEKSLFQEGSRAFFPACYSGDGRLKSWGLTLRCALVVFTLLM 446
mVIAAT 374 LPG--SIRA----VVNLFLVAKALLSYPLPFFAAVEV---LEKSLFQEGSRAFFPACYGGDGRLKSWGLTLRCALVVFTLLM 446
xVIAAT 367 LPS--TIRA----VVNLFLVAKALLSYPLPFFAAVEV---LEKSLFQEGARAFFPNCYGGDGRLKSWGLTLRCALVVFTLLM 439
zVIAAT 379 LPS--SIRA----VVNLFLVSKALLSYPLPFFAAVEV---LEKTFFQDGGRAFFPDCYGGDGRLKSWGLSLRCALVVFTMLM 451
dVIAAT 397 LHSQ-GFKG----MVNFFLVIKALLSYPLPYYAACEL---LERNFFRGPPKTKFPTIWNLDGELKVWGLGFRVGVIVSTILM 470
UNC-47 335 LPNQ-SFKI----LVNLILVVKALLSYPLPFYAAVQL---LKNNLFLGYPQTPFTSCYSPDKSLREWAVTLRIILVLFTLFV 408
TM10
TM11
hVIAAT 447 AIYVPHFALLMGLTGSLTGAGLCFLLPSLFHLRLLWRKLLWHQVFFDVAIFVIGGICSVSGFVHSLEGLIEAYRTNAED 525
mVIAAT 447 AIYVPHFALLMGLTGSLTGAGLCFLLPSLFHLRLLWRKLLWHQVFFDVAIFVIGGICSVSGFVHSLEGLIEAYRTNAED 525
xVIAAT 440 AIYVPHFALLMGLTGSLTGAGLCFLLPSLFHLKLLWRKLQWHQVFFDVSIFVIGSICSVSGFVHSLEGLIEAFRFNIED 518
zVIAAT 452 AIYVPHFALLMGLTGSLTGAGLCFLLPSLFHLKLLWRKLLWHQVFFDVAIFVIGGICSISGFIHSVEGLIEAYKYNLPD 530
dVIAAT 471 AIFIPHFSILMGFIGSFTGTMLSFIWPCYFHIKIKGHLLDQKEIAKDYLIIGLGVLFGVIGIYDSGNALINAFEIGLPF 549
UNC-47 409 ALSVPYLVELMGLVGNITGTMLSFIWPALFHLYIKEKTLNNFEKRFDQGIIIMGCSVCISGVYFSSMELLRAINSADS- 486
Key residues involved in: Substrate binding Chloride ion binding Protonation Conformational transition

### Extended Data Figure 11. Sequence alignment of VIAAT from different species.

Secondary structural elements of human VIAAT (hVIAAT; UniProt: Q9H598) are shown above the sequence alignment, with unmodeled loop regions indicated by dashed lines. Sequences of mouse VIAAT (mVIAAT; UniProt: O35633), *Xenopus laevis* VIAAT (xVIAAT; UniProt: Q6DIV6), zebrafish VIAAT (zVIAAT; UniProt: A1L1T3), *Drosophila* VIAAT (dVIAAT; UniProt: A1Z9M6), and *Caenorhabditis elegans* VIAAT (UNC-47; UniProt: P34579) were aligned to human VIAAT using Clustal Omega and visualized with Jalview. Conserved residues are colored red (acidic), blue (basic), or gray (other). Residues involved in substrate binding, chloride coordination, proton coupling, and conformational transition are marked with orange triangles, green circles, purple pentagons, and cyan square, respectively.

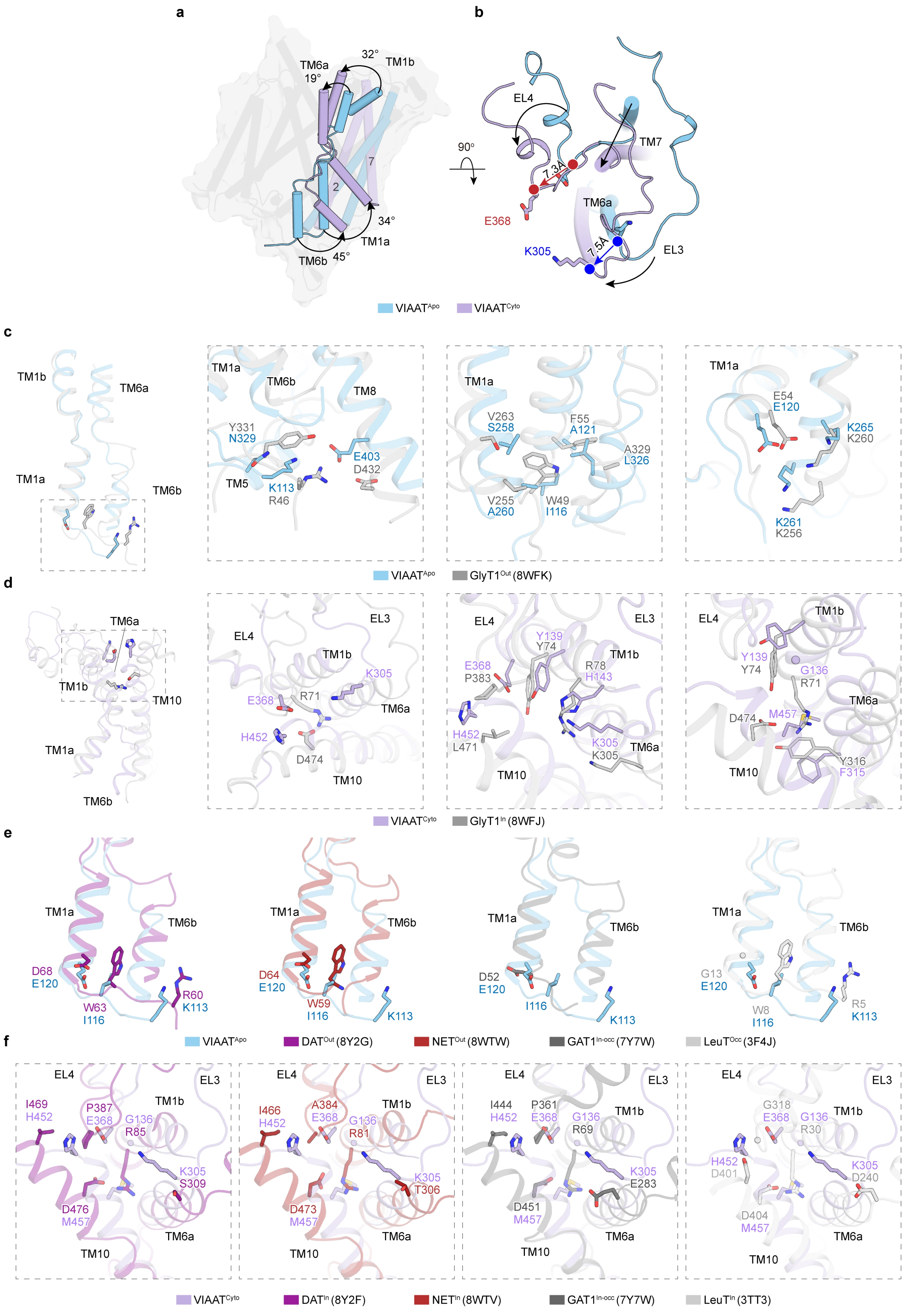

a
b
TM6a
19°
32°
TM1b
2
7
34°
TM1a
TM6b
45°
EL4
90°
7.3Å
TM7
E368
TM6a
7.5Å
K305
EL3
VIAAT^Apo
VIAAT^Cyto
c
TM1b
TM6a
TM1a
TM6b
TM1a
TM6b
TM8
Y331
N329
E403
D432
TM5
K113
R46
TM1a
V263
S258
F55
A121
A329
L326
V255
A260
W49
I116
TM1a
E54
E120
K265
K260
K261
K256
VIAAT^Apo
GlyT1^Out (8WFK)
d
TM6a
TM1b
TM10
TM1a
TM6b
EL4
EL3
TM1b
K305
E368
R71
TM6a
H452
D474
TM10
EL4
EL3
Y139
Y74
E368
P383
R78
H143
TM1b
H452
L471
K305
K305
TM6a
TM10
EL4
TM1b
Y139
Y74
G136
R71
TM6a
D474
M457
TM10
Y316
F315
VIAAT^Cyto
GlyT1^In (8WFJ)
e
TM1a
TM6b
D68
E120
W63
I116
R60
K113
TM1a
TM6b
D64
E120
W59
I116
K113
TM1a
TM6b
D52
E120
I116
K113
TM1a
TM6b
G13
E120
W8
I116
R5
K113
f
VIAAT^Apo
DAT^Out (8Y2G)
NET^Out (8WTW)
GAT1^In-occ (7Y7W)
LeuT^Occ (3F4J)
EL4
EL3
I469
H452
P387
E368
G136
R85
TM1b
K305
S309
D476
M457
TM6a
TM10
EL4
EL3
I466
H452
A384
E368
G136
R81
TM1b
K305
T306
D473
M457
TM6a
TM10
EL4
EL3
I444
H452
P361
E368
G136
R69
TM1b
K305
E283
D451
M457
TM6a
TM10
EL4
EL3
G318
E368
G136
R30
TM1b
H452
D401
K305
D240
D404
M457
TM6a
TM10
VIAAT^Cyto
DAT^In (8Y2F)
NET^In (8WTV)
GAT1^In-occ (7Y7W)
LeuT^In (3TT3)

### Extended Data Figure 12. Distinct structural elements contribute to conformational transitions in VIAAT compared with neurotransmitter sodium symporters.

**a, b,** Side view (a) and top-down view (b) of the structural comparison between $VIAAT^{Apo}$ (cyan) and cytosol-facing $VIAAT^{Cyto}$ (purple), aligned based on the scaffold domain, highlighting conformational differences.

**c,** Comparison of interactions stabilizing the lumen-facing conformation of $VIAAT^{Apo}$ (cyan) and the outward-facing $GlyT1^{SSR}$ structure (PDB: 8WFK; grey). Key residues involved in interactions are shown as sticks.

**d,** Comparison of interactions stabilizing the cytosol-facing conformation of VIAAT and the inward-facing $GlyT1^{ALX}$ structure (PDB: 8WFJ). Key residues involved in interactions are depicted as sticks.

**e,** Structural comparison of interaction networks stabilizing the outward conformation in SLC6 transporters, including DAT (PDB: 8Y2G), NET (PDB: 8WTW), GAT1 (PDB: 7Y7W), and LeuT (PDB: 3F4J), and the corresponding residues in VIAAT. Key residues are shown as sticks.

**f,** Structural comparison of interaction networks stabilizing the inward conformation in SLC6 transporters, including DAT (PDB: 8Y2F), NET (PDB: 8WTV), GAT1 (PDB: 7Y7W), and LeuT (PDB: 3TT3), and the corresponding residues in VIAAT. Key residues are shown as sticks.

Extended Data Table 1. Cryo-EM data collection, refinement and validation statistics.

| | VIAAT$^{Apo}$ (EMD-67826) (PDB: 21MN) | VIAAT$^{GABA}$ (EMD-67799) (PDB: 21LK) | VIAAT$^{Gly}$ (EMD-67798) (PDB: 21LJ) | VIAAT$^{PO4}$ (EMD-67797) (PDB: 21LI) |
|---|---|---|---|---|
| **Data collection and processing** | | | | |
| Magnification | 105,000 × | 105,000 × | 105,000 × | 105,000 × |
| Voltage (kV) | 300 | 300 | 300 | 300 |
| Electron exposure ($e^{-}$/Å$^2$) | 60 | 60 | 60 | 60 |
| Defocus range (μm) | -1.2 ~ -2.2 | -1.2 ~ -2.2 | -1.2 ~ -2.2 | -1.2 ~ -2.2 |
| Pixel size (Å) | 0.85 | 0.85 | 0.85 | 0.85 |
| Symmetry imposed | C1 | C1 | C1 | C1 |
| Initial particle images (no.) | 2,335,331 | 3,191,330 | 4,759,153 | 1,858,894 |
| Final particle images (no.) | 207,791 | 88,339 | 243,502 | 54,346 |
| Map resolution (Å) | 3.3 | 2.7 | 3.1 | 3.0 |
| FSC threshold | 0.143 | 0.143 | 0.143 | 0.143 |
| **Refinement** | | | | |
| Initial model used (PDB code) | | | | |
| Model resolution (Å) | 3.4 | 3.0 | 3.3 | 3.2 |
| FSC threshold | 0.5 | 0.5 | 0.5 | 0.5 |
| Map sharpening *B* factor (Å$^2$) | 131.7 | 90.2 | 136.0 | 97.7 |
| Model composition | | | | |
| Non-hydrogen atoms | 3,051 | 3,131 | 3,053 | 2,789 |
| Protein residues | 391 | 398 | 389 | 389 |
| Ligands | 0 | 1 | 1 | 0 |
| *B* factors (Å$^2$) | | | | |
| Protein | 35.77 | 35.46 | 47.36 | 39.02 |
| Ligand | 0 | 50.75 | 63.83 | 0 |
| R.m.s.d deviations | | | | |
| Bond lengths (Å) | 0.005 | 0.003 | 0.007 | 0.004 |
| Bond angles (°) | 0.775 | 0.632 | 1.074 | 0.602 |
| Validation | | | | |
| MolProbity score | 1.83 | 1.37 | 1.52 | 1.45 |
| Clash score | 5.00 | 3.01 | 5.18 | 4.22 |
| Ramachandran plot | | | | |
| Favored (%) | 94.57 | 95.94 | 96.36 | 96.28 |
| Allowed (%) | 5.43 | 4.06 | 3.64 | 3.72 |
| Disallowed (%) | 0 | 0 | 0 | 0 |

Extended Data Table 2. Epilepsy-associated missense mutations in VIAAT.

| **Mutations** | **Position** |
|---|---|
| G43C | IL1 |
| A90T | IL1 |
| V263A/M | TM5 |
| L269P | TM5 |
| F322C | TM6 |
| M330T | IL4 |
| P395L | TM8 |
| L445F | TM9 |
| G461D | TM10 |
| T464R | TM10 |
| G465S | TM10 |
| L468P | TM10 |
